# ppGpp regulates transcription elongation via direct and indirect inputs to RNA polymerase pausing and nucleotide addition

**DOI:** 10.64898/2026.05.13.724835

**Authors:** Andreas U. Mueller, Rachel A. Mooney, Michael D. Engstrom, Yu Bao, Michael B. Wolfe, Balendra Sah, Jonathan Buscher, Jason Saba, James Liu, Seth A. Darst, Robert Landick

## Abstract

The signaling molecules guanosine 5′-tri/diphosphate 3′-diphosphate, (p)ppGpp, control bacterial protein synthesis rates and cell growth by targeting transcription, translation, NTP synthesis, and other functions. In lineages like *E. coli*, (p)ppGpp produced in response to charged-tRNA deficiency directly targets transcribing RNAP polymerase (RNAP) to match its pace to the pioneering ribosome on the nascent RNA (transcription–translation coupling). However, the mechanism by which (p)ppGpp slows RNAP is poorly defined. (p)ppGpp may allosterically stimulate RNAP pausing, inhibit catalysis, promote backtracking, compete for substrate GTP, inhibit GTP synthesis, or uncouple transcription–translation by inhibiting translation. Using a combination of cryo-EM, biochemical assays, and quantitative nascent elongating transcript sequencing (qNET-seq), we establish that (p)ppGpp allosterically regulates pausing and nucleotide addition via distinct motions of the RNAP swivel module and both competes with and lowers GTP in vivo. (p)ppGpp stimulates swiveling at pause sites to delay escape but may also inhibit counter-swiveling required in every round of nucleotide addition.

**Highlights:**

- ppGpp biases RNAP toward swiveling and away from a catalytically-competent state
- ppGpp effects on RNAP conformation explain ppGpp stimulation of RNAP pausing
- ppGpp stimulation of RNAP pausing in vivo is mediated mainly by reduced GTP levels
- ppGpp may allosterically slow RNAP and increases gene occupancy in vivo

## INTRODUCTION

The bacterial alarmone (p)ppGpp links bacterial cell growth to nutrient availability, especially amino-acid availability.^1^ Diverse targets, functions, and routes of (p)ppGpp synthesis exist, but a key route depends on the enzyme RelA, activated by association with uncharged tRNA–bound ribosomes (Figure 1A).^2^ (p)ppGpp levels directly or indirectly modulate transcription initiation and elongation,^3–5^ translation initiation and elongation,^6^ DNA replication and repair,^7,8^ and NTP synthesis and salvage.^9^ Amino-acid starvation and other stresses trigger a stringent response in which mM levels of (p)ppGpp dramatically reconfigure cell processes.^10,11^ During normal cell growth, lower but varying (p)ppGpp levels link transcription and translation rates to cell growth rate. In *E. coli*, most (p)ppGpp is present as ppGpp (used for simplicity hereafter).^12^

**Figure 1.**
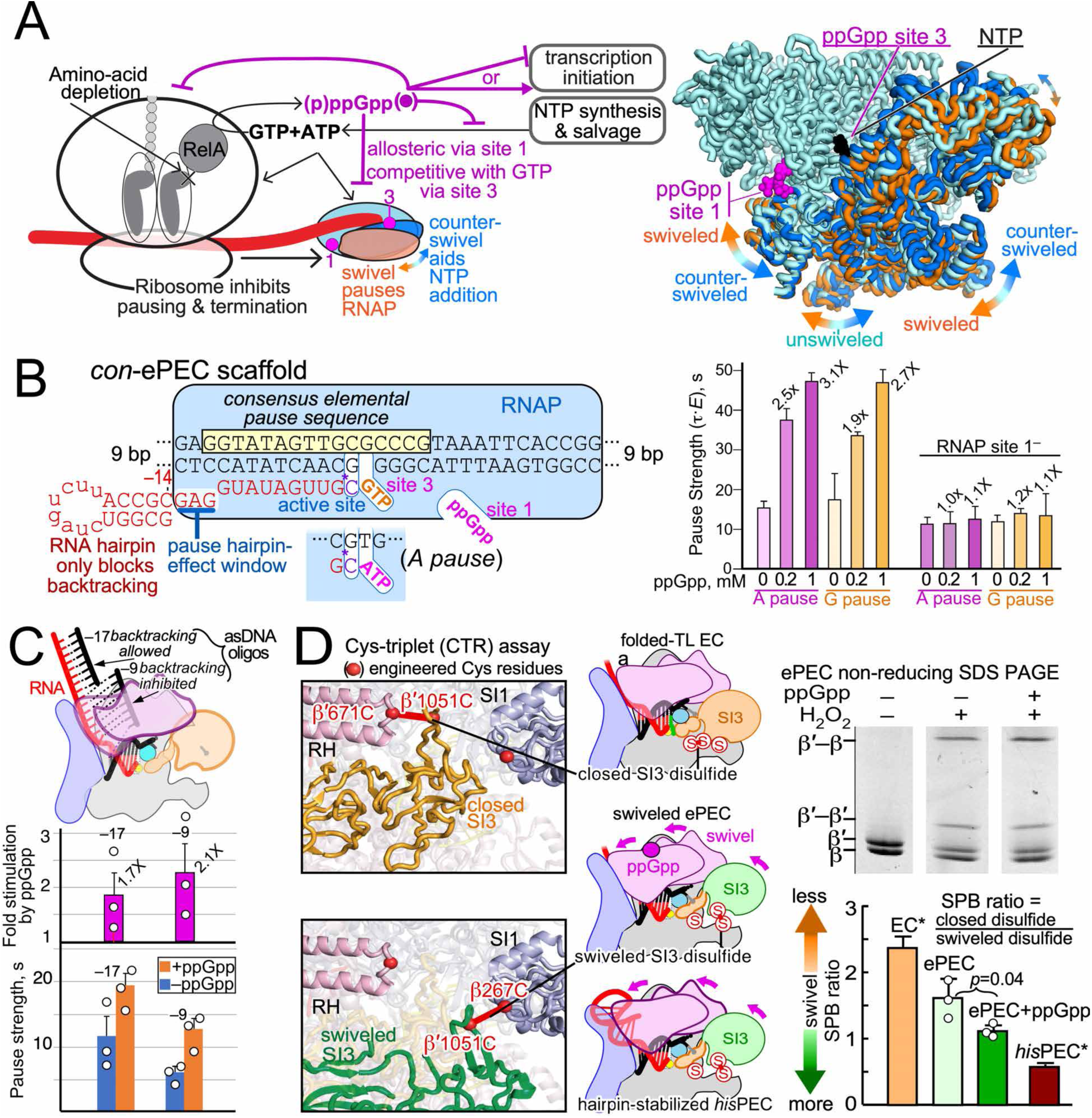
ppGpp binding to RNAP favors RNAP swiveling. (A) ppGpp-mediated regulation in *E. coli*. ppGpp (magenta) is produced when uncharged tRNA in ribosomes activate RelA. ppGpp inhibits translation factors (IF-2, EF-G, EF-Tu), RNAP initiation and elongation, and NTP synthesis. RNAP elongation is aided when ribosomes remain coupled to RNAP. ppGpp binds RNAP site 1 near the swivel module, which rotates up to ∼7° between swiveled (orange) and counter-swiveled (blue) positions. (B) ppGpp (0.2 or 1 mM) stimulates pausing allosterically before either A or G addition on a consensus elemental pause scaffold, but not when site 1 is eliminated (Figure S1B). (C) Antisense DNAs (asDNAs) demonstrate that ppGpp enhances pausing independent of backtracking. A backtracking ePEC was created by active elongation of promoter-initiated ECs. Pause strengths (ρ) without (blue bars) or with (orange bars) ppGpp (100 µM) revealed a similar fold stimulation by ppGpp (magenta bars) whether asDNAs allowed (–17 asDNA) or inhibited (–9 asDNA) backtracking (Figures S1C and S1D). (D) The Cys-triplet (CTR) assay reveals ppGpp stimulates swiveling of ePEC. RNAP diagrams show Cys positions in an EC with bound substrate NTP (top) and in swiveled PECs without and with a pause hairpin (middle and bottom).^67^ The CTR assay measures the swivel positional bias (SPB) ratio using differential migration of CTR crosslinked RNAP subunits during non-reducing SDS-PAGE (gel panels). SPB of an active EC, the ePEC used for cryo-EM without or with ppGpp, and the hairpin-stabilized *his*PEC revealed that ppGpp biases RNAP toward swiveling. Results for the EC and *his*PEC (asterisks) were published previously.^67^

Although ppGpp action mechanisms for many targets are now defined, how ppGpp adjusts RNAP elongation rates remains poorly understood. In some bacteria (*e.g*., proteobacteria like *E. coli*), ppGpp directly binds RNA polymerase (RNAP) at three sites.^13,14^ Allosteric site 1 at the interface of the μ and β′ subunits primarily affects elongation. Allosteric site 2 at the interface of the NTP-entry channel and dissociable DksA only affects initiation. Site 3 is the RNAP active site, where >∼0.5 mM ppGpp competes with substrate GTP. ppGpp binds the allosteric sites 1 and 2 in *Eco*RNAP with roughly equal *K*_D_^app^ of ∼6 µM^15^ whereas it slows RNA chain extension and stimulates pausing by *Eco*RNAP noncompetitively (*i.e*., allosterically via site 1) with a *K*_i_ of ∼50 µM in vitro.^16^

Pausing is an evolved feature of RNAPs that punctuates RNA chain elongation in response to DNA sequence, RNA structure, and transcription factors to control the rate of RNA synthesis and enable its regulation.^17,18^ Pausing occurs by an initial, sequence-dependent isomerization of an elongation complex (EC) into an elemental paused EC (ePEC).^17,19–21^ The ePEC enters more long-lived pause states by backtracking (reverse translocation) of RNA and DNA through RNAP, by binding of regulators, or by formation of a pause RNA hairpin in the RNA exit channel of RNAP. The ePEC itself can assume multiple conformations: (1) pre-translocated, with a closed active site and folded TL (resembling NTP-bound ECs); (2) pre-translocated and semi-closed (TL folded with open rim helices and F-loop); (3) pre-translocated and open (TL unfolded); (4) half-translocated (RNA but not DNA translocated); and (5) swiveled.^22^ RNAP swiveling involves a concerted ∼5° rotation of the RNAP swivel module (RNAP shelf, clamp, dock, jaw, SI3, β′ C-terminal segment, and μ) compared to the EC conformation (e.g., PDB 6ALH)^23^ and an associated bend in the bridge helix (BH).^19,20,24,25^ In a swiveled ePEC, the bent BH inhibits both DNA translocation and NTP binding and the rotated SI3 inhibits folding of the trigger loop (TL) required for catalysis (Figures 1A and S1A). The swivel module rotates in the opposite direction by ∼1.5° (counter-swivels) in the NTP-bound, closed EC.^26^

Contacts of ppGpp with μ and β′ side chains in site 1 of a non-paused *Eco*RNAP EC are well defined by cryo-EM, but cause no significant conformational changes relative to the unbound EC.^27^ Nonetheless, an RNAP conformational change is proposed to be responsible for ppGpp–site 1 activity.^16,28^ An attractive possibility is that ppGpp may modulate RNAP swiveling, since site 1 lies at the interface of the swivel and core RNAP modules (Figure 1A).^28^ Both nascent RNA structures and transcription factors like NusA and NusG from some bacteria (*e.g*., *M. tuberculosis*) can stimulate RNAP pausing by stabilizing swiveled states,^19,20,25^ whereas coupled ribosomes in *E. coli* and NusGs from other lineages (*e.g*., *E. coli*) inhibit swiveling and pausing.^25,29,30^ Alternatively, ppGpp may promote paused EC (PEC) backtracking.^8^

Elevated ppGpp slows RNAP in vivo,^31–33^ either via allosteric or competitive effects on RNAP or via ppGpp-induced reductions in translation or NTP levels. ppGpp reduces translation and NTP levels both directly by binding relevant enzymes (e.g., IF-2, EF-G, EF-Tu, PurF for GTP biosynthesis, and Gsk for GTP salvage)^9,34,35^ and indirectly by altering gene transcription.^12,36^ Translation and NTP levels affect RNAP transcript elongation and pausing via transcription–translation coupling^37^ and as substrates for RNA synthesis, respectively. These effects are difficult to disentangle due to complex regulation.

To elucidate how ppGpp controls pausing by *Eco*RNAP, we combined multiple biochemical approaches with cryo-EM and quantitative NET-seq and ChIP-seq (qNET-seq and qChIP-seq) analysis of transcription in vivo. Use of elemental PECs allowed detection of ppGpp effects on swiveling. qNET-seq, which uses a ‘spike-in’ reference, enabled genome-wide quantitation of RNAP occupancy on DNA at single-bp resolution and dissection of the direct versus indirect effects of ppGpp.

## RESULTS

### ppGpp directly targets the elemental PEC via site 1

To enable a structural elucidation of the effect of ppGpp on pausing, we sought to define a discrete PEC whose activity is allosterically modulated by ppGpp via site 1. Although pausing by *Eco*RNAP is prominent in vivo before addition of G (G pauses),^38–40^ *Eco*RNAP pauses before A and G similarly in vitro.^39,41^ Thus, to cleanly distinguish effects of ppGpp binding to site 1 versus in the RNAP active site (site 3), we compared ‘G pause’ and ‘A pause’ versions of the *E. coli* consensus elemental pause site (Figure 1B). At 200 µM, ppGpp increased pause strength for the G and A consensus elemental PECs (*con*-ePECs) by 1.9- and 2.5-fold, respectively (Figure 1B, Table S1). Increasing ppGpp to 1 mM increased pause strength further. ppGpp even at 1 mM had no effect on the A pause when site 1 was eliminated in *Eco*RNAP by substitutions shown previously to abolish ppGpp binding.^12,28,42^ A slight residual effect of ppGpp on the G pause with RNAP site 1^−^ may reflect competitive inhibition of GTP binding in site 3. We conclude that the ppGpp allosteric effect on *Eco*RNAP can be isolated using the ‘A’ *con*-ePEC scaffold.

### ppGpp stimulates RNAP pausing by enhancing RNAP swiveling not backtracking

ppGpp may promote backtracking,^8^ a possible explanation for its effect on pausing but also a potential complication to structural analysis of the ppGpp allosteric effect on RNAP. To ask if backtracking is required for ppGpp stimulation of pausing, we measured the ppGpp effect on the *con*-ePEC on a template prone to backtracking (Figures S1C and S1D). Using antisense DNA (asDNA) oligos that can block backtracking by annealing to the nascent RNA, we assessed the role of backtracking in ppGpp action by progressively advancing the asDNA toward the pause RNA 3′ end. ppGpp increased pause strength ∼2-fold regardless of asDNA location and effect of the asDNA on pausing (Figures 1C and S1D, Table S1). asDNA affected pausing in three distinct phases. From –17 to –14 (phase 1), pause strength decreased, consistent with inhibition of backtracking.^43^ From –13 to –11 (phase 2), asDNA pairing can weakly mimic the action of a PH^44^ and thus increased pausing even though backtracking is inhibited. At –10 and –9 (phase 3), the PH-like effect is lost due to changes in duplex contacts.^45,46^ ppGpp-stimulation of pausing was unaffected whether backtracking was allowed or blocked, consistent with ppGpp directly affecting the non-backtracked ePEC. We conclude that ppGpp stimulation of pausing does not require backtracking but rather operates directly on the *con*-ePEC in its initial register.

Based on these results, we predicted that addition of ppGpp to the *con*-ePEC blocked for backtracking should reveal changes in RNAP responsible for ppGpp action. To block backtracking that is known to occur in the *con-*PEC and that would complicate cryo-EM analysis,^21^ we positioned a nascent RNA hairpin just outside the window within which hairpins can increase pausing (RNA 3′ end–proximal bp in hairpin at –14; Figure S1A). We then used a Cys-triplet (CTR) disulfide crosslinking assay to assess the effect of ppGpp on RNAP swiveling in the backtrack-blocked *con*-ePEC. In the CTR assay, a site-specifically located Cys on SI3 (β′D1051C) forms a disulfide with βR267C in SI1 in a swiveled RNAP and instead forms a disulfide with β′G671C in the rim helices in a counter-swiveled, TL-folded RNAP (Figure 1D, Table S1).^47^ When ppGpp was added to the backtrack-blocked ePEC, crosslinking shifted toward the swiveled conformation (Fig. 1D). We conclude that in vitro, ppGpp can stimulate pausing allosterically via site 1 by stimulating RNAP swiveling.

### ppGpp shifts cryo-EM PEC populations toward swiveled states

To test for ppGpp effects on RNAP conformation and swiveling directly, we next used cryo-EM to characterize the conformational distributions of ePECs with and without ppGpp (Figures 2, S2, S3, and S4; Table 1). We obtained consensus reconstructions at global resolutions of 2.5 Å and 2.8 Å for the −ppGpp and +ppGpp conditions, respectively. After removal of particles containing σ^70^ or lacking ω, conformational heterogeneity was resolved by principal component analysis. We found that *Eco*RNAP reconstituted on the A-pause *con*-ePEC scaffold into the full suite of previously defined ePEC conformations (states *a*–*e*; Figures 2A and 2B).^22^ However, state *d* (partially swiveled ePEC) was a heterogeneous mixture of partially swiveled states and thus was not modeled with atomic coordinates (Figure S2). ppGpp shifted the distribution of conformations away from TL-folded states toward TL-unfolded states and away from counter-swiveled, unswiveled, and partially swiveled states toward the fully swiveled state (Figures 2C, 2D, and 2E). Notably, the heterogeneous partially swiveled states (state *d*) disappeared in the presence of ppGpp. The relatively small, ppGpp-induced shift in the distribution of interconverting, energetically similar ePEC states suggests a relatively subtle energetic effect of ppGpp consistent with the CTR assay results (Figure 1D). To assess swiveling fully, we measured the rotation angles of the swivel module.^19,20,22,30^ Notably, the swivel module was counter-swiveled in the closed ePEC (–2.32° rotation), as also observed in NTP-bound, closed ECs (Figures 2B, S4B, and S4C; Table 1).^26^ Further, for every state except the fully swiveled ePEC, binding of ppGpp slightly shifted the swivel module toward swiveling (Table 1; Figures S4B and S4C). We conclude that ppGpp acts on the elemental pause by modestly stabilizing the fully swiveled state (state *e*) and modestly destabilizing the counter-swiveled, closed ePEC state (state *a*).

**Figure 2.**
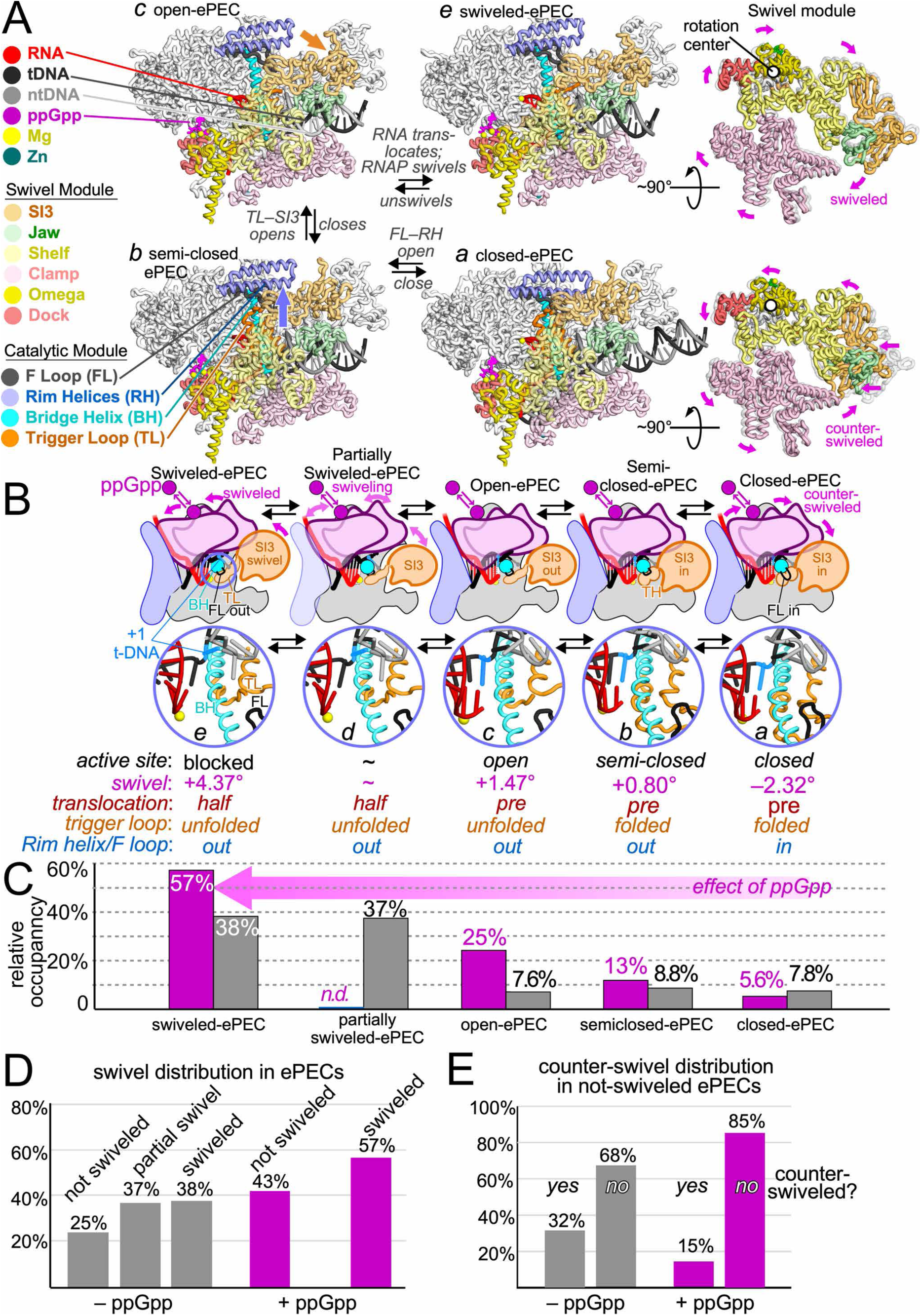
cryo-EM structures of ppGpp-bound paused ePEC. (A) Conformation of RNAP in ePEC structures. Cα backbone depictions of RNAP with the different parts of the swivel module in pale colors and the catalytic module consisting of the bridge helix (BH, cyan), trigger loop (TL, orange), F-loop (FL, dark gray), and rim helices (RH, blue) highlighted (*a*–*e* refer to states as lettered in Kang et al., 2023 and as shown in (B)).^22^ *a*, pretranslocated ePEC with folded TL and FL–RH in the down/in position; *b*, pretranslocated ePEC with folded TL and FL–RH in the up/out position; *c*, pretranslocated ePEC with unfolded TL; and *e*, the swiveled ePEC. The partially swiveled ePEC states (*d*) were not modeled as a discrete state. (B) ePEC states *a*–*e* depicted as RNAP cartoons. The insets highlight Cα and nucleic acid cartoons of the active-site configurations for each state. Key features of each state are shown below the insets. (C) The observed distribution of ePEC particles classified into the different ePEC states in the absence (gray bars) and presence (magenta bars) of bound ppGpp. The magenta arrow highlights the ppGpp-induced shift in ePEC state distribution away from state *a* and toward the swiveled state *e*. (D) The proportions of ePECs with in unswiveled, partially swiveled, and fully swiveled states in the absence (gray) and presence (magenta) of ppGpp. (E) The proportions of ePECs in the counter-swiveled state in the absence (gray) and presence (magenta) of ppGpp.

**Table 1:**
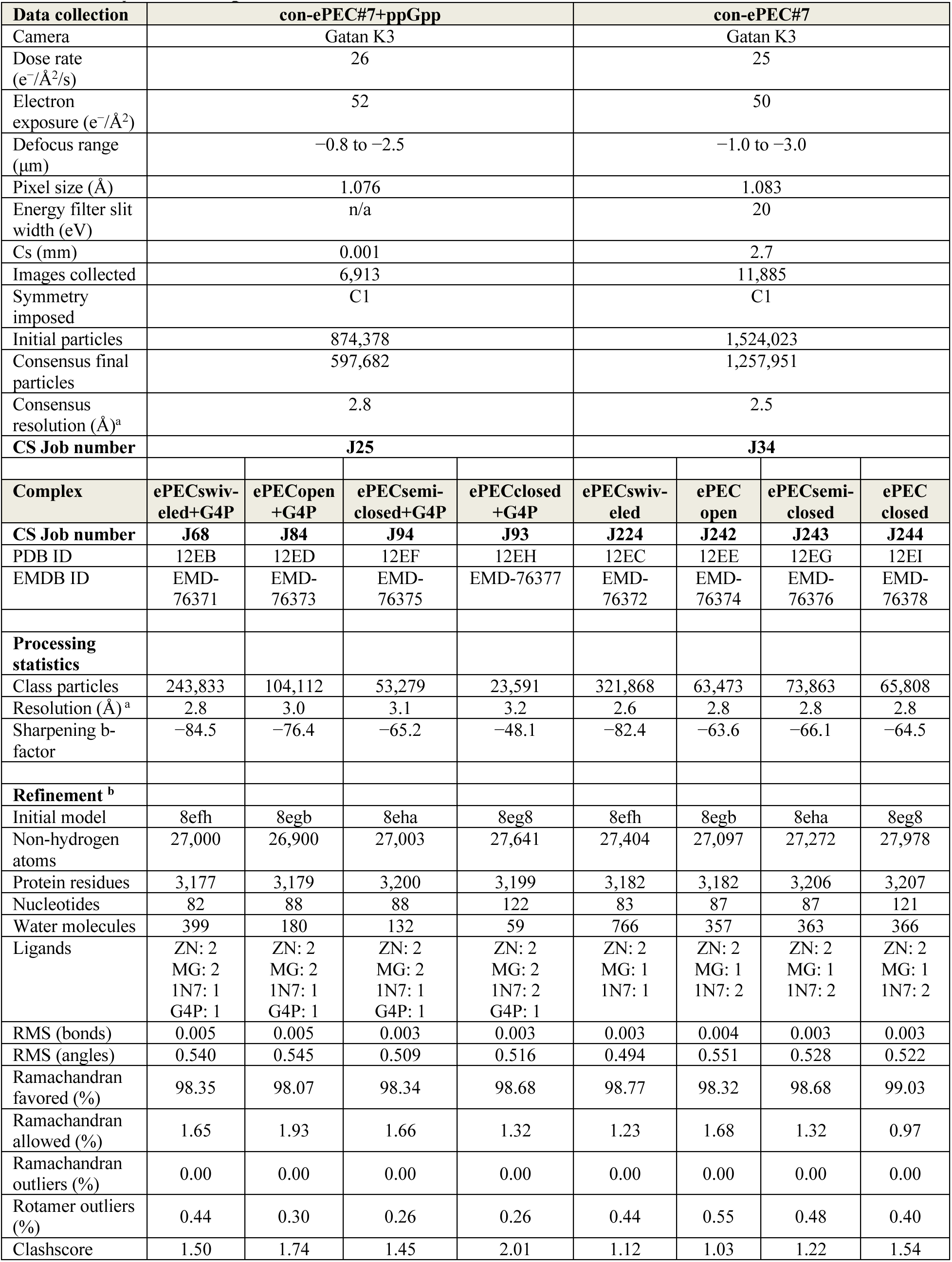
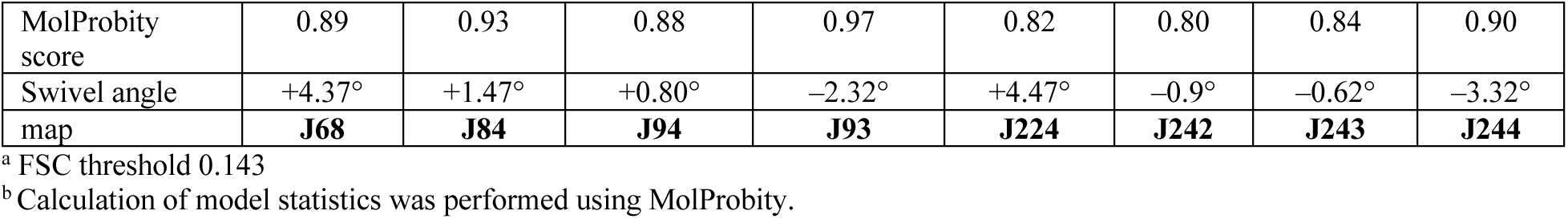
Cryo-EM map and model statistics.

| Data collection | con-ePEC#7+ppGpp |  |  |  | con-ePEC#7 |  |  |  |
| --- | --- | --- | --- | --- | --- | --- | --- | --- |
| Camera | Gatan K3 |  |  |  | Gatan K3 |  |  |  |
| Dose rate<br>(e<sup>-</sup>/Å<sup>2</sup>/s) | 26 |  |  |  | 25 |  |  |  |
| Electron<br>exposure (e<sup>-</sup>/Å<sup>2</sup>) | 52 |  |  |  | 50 |  |  |  |
| Defocus range<br>(μm) | -0.8 to -2.5 |  |  |  | -1.0 to -3.0 |  |  |  |
| Pixel size (Å) | 1.076 |  |  |  | 1.083 |  |  |  |
| Energy filter slit<br>width (eV) | n/a |  |  |  | 20 |  |  |  |
| Cs (mm) | 0.001 |  |  |  | 2.7 |  |  |  |
| Images collected | 6,913 |  |  |  | 11,885 |  |  |  |
| Symmetry<br>imposed | C1 |  |  |  | C1 |  |  |  |
| Initial particles | 874,378 |  |  |  | 1,524,023 |  |  |  |
| Consensus final<br>particles | 597,682 |  |  |  | 1,257,951 |  |  |  |
| Consensus<br>resolution (Å)<sup>a</sup> | 2.8 |  |  |  | 2.5 |  |  |  |
| CS Job number | J25 |  |  |  | J34 |  |  |  |
| Complex | ePECswiv-<br>eled+G4P | ePECopen<br>+G4P | ePECsemi-closed+G4P | ePECclosed<br>+G4P | ePECswiv-<br>eled | ePEC<br>open | ePECsemi-closed | ePEC<br>closed |
| CS Job number | J68 | J84 | J94 | J93 | J224 | J242 | J243 | J244 |
| PDB ID | 12EB | 12ED | 12EF | 12EH | 12EC | 12EE | 12EG | 12EI |
| EMDB ID | EMD-76371 | EMD-76373 | EMD-76375 | EMD-76377 | EMD-76372 | EMD-76374 | EMD-76376 | EMD-76378 |
| Processing<br>statistics |  |  |  |  |  |  |  |  |
| Class particles | 243,833 | 104,112 | 53,279 | 23,591 | 321,868 | 63,473 | 73,863 | 65,808 |
| Resolution (Å)<sup>a</sup> | 2.8 | 3.0 | 3.1 | 3.2 | 2.6 | 2.8 | 2.8 | 2.8 |
| Sharpening b-factor | -84.5 | -76.4 | -65.2 | -48.1 | -82.4 | -63.6 | -66.1 | -64.5 |
| Refinement<sup>b</sup> |  |  |  |  |  |  |  |  |
| Initial model | 8efh | 8egb | 8eha | 8eg8 | 8efh | 8egb | 8eha | 8eg8 |
| Non-hydrogen<br>atoms | 27,000 | 26,900 | 27,003 | 27,641 | 27,404 | 27,097 | 27,272 | 27,978 |
| Protein residues | 3,177 | 3,179 | 3,200 | 3,199 | 3,182 | 3,182 | 3,206 | 3,207 |
| Nucleotides | 82 | 88 | 88 | 122 | 83 | 87 | 87 | 121 |
| Water molecules | 399 | 180 | 132 | 59 | 766 | 357 | 363 | 366 |
| Ligands | ZN: 2<br>MG: 2<br>1N7: 1<br>G4P: 1 | ZN: 2<br>MG: 2<br>1N7: 1<br>G4P: 1 | ZN: 2<br>MG: 2<br>1N7: 1<br>G4P: 1 | ZN: 2<br>MG: 2<br>1N7: 2<br>G4P: 1 | ZN: 2<br>MG: 1<br>1N7: 1 | ZN: 2<br>MG: 1<br>1N7: 2 | ZN: 2<br>MG: 1<br>1N7: 2 | ZN: 2<br>MG: 1<br>1N7: 2 |
| RMS (bonds) | 0.005 | 0.005 | 0.003 | 0.003 | 0.003 | 0.004 | 0.003 | 0.003 |
| RMS (angles) | 0.540 | 0.545 | 0.509 | 0.516 | 0.494 | 0.551 | 0.528 | 0.522 |
| Ramachandran<br>favored (%) | 98.35 | 98.07 | 98.34 | 98.68 | 98.77 | 98.32 | 98.68 | 99.03 |
| Ramachandran<br>allowed (%) | 1.65 | 1.93 | 1.66 | 1.32 | 1.23 | 1.68 | 1.32 | 0.97 |
| Ramachandran<br>outliers (%) | 0.00 | 0.00 | 0.00 | 0.00 | 0.00 | 0.00 | 0.00 | 0.00 |
| Rotamer outliers<br>(%) | 0.44 | 0.30 | 0.26 | 0.26 | 0.44 | 0.55 | 0.48 | 0.40 |
| Clashscore | 1.50 | 1.74 | 1.45 | 2.01 | 1.12 | 1.03 | 1.22 | 1.54 |
| MolProbity score | 0.89 | 0.93 | 0.88 | 0.97 | 0.82 | 0.80 | 0.84 | 0.90 |
| Swivel angle | +4.37° | +1.47° | +0.80° | −2.32° | +4.47° | −0.9° | −0.62° | −3.32° |
| map | **J68** | **J84** | **J94** | **J93** | **J224** | **J242** | **J243** | **J244** |

<sup>a</sup> FSC threshold 0.143
<sup>b</sup> Calculation of model statistics was performed using MolProbity.

### ppGpp contacts are altered in the counterswiveled ePEC

To understand how ppGpp affects RNAP, we examined the ppGpp-bound site 1 in the ePEC states. ppGpp contacts in the swiveled ePEC (at global resolution 2.6 Å; estimated local resolution of 2.8 Å at site 1; Figure S3A) were largely indistinguishable from those reported for a stable EC (Figures 3A–3C).^31^ The guanine ring is positioned by π-Arg362 stacking opposite a hydrophobic contact to Ile619 and N2–Asp662 and O6–Gln623 hydrogen bonds involving relatively immobile parts of RNAP, including the double-λλ′ β barrel module that anchors the catalytic Mg^2+^ in the RNAP active site. The β′ core α-helix (614-635) H-bonds to the ribose sugar and 3′ diphosphate (Lys615 to 2′-O, 3′-O, and a 3′-βP-O). However, most 5′ diphosphate contacts are made by parts of the swivel module: the N-terminal residues of the μ subunit (Ala2 and Arg3) and a key salt bridge between the 5′-β phosphate Arg417 in the dock. Although the stable EC is not swiveled, these phosphate contacts are largely unchanged in the swiveled ePEC (Figure 3). However, changes in ppGpp position, sugar–diphosphate conformation, and ppGpp contacts were evident when the dock and μ moved away from ppGpp in the counter-swiveled, closed ePEC (Figures 3D–3F). The guanine ring and ribose shifted ∼1 Å away from the core ppGpp binding pocket and the sugar–diphosphate twisted, resulting in apparent loss of interaction with Arg317. Cryo-EM signal for the ppGpp metal ions was reduced, although the lower overall resolution of the active site–closed ePEC limited our ability to determine changes in site-1 conformation and occupancy (3.2 Å from ∼24K particles).

**Figure 3.**
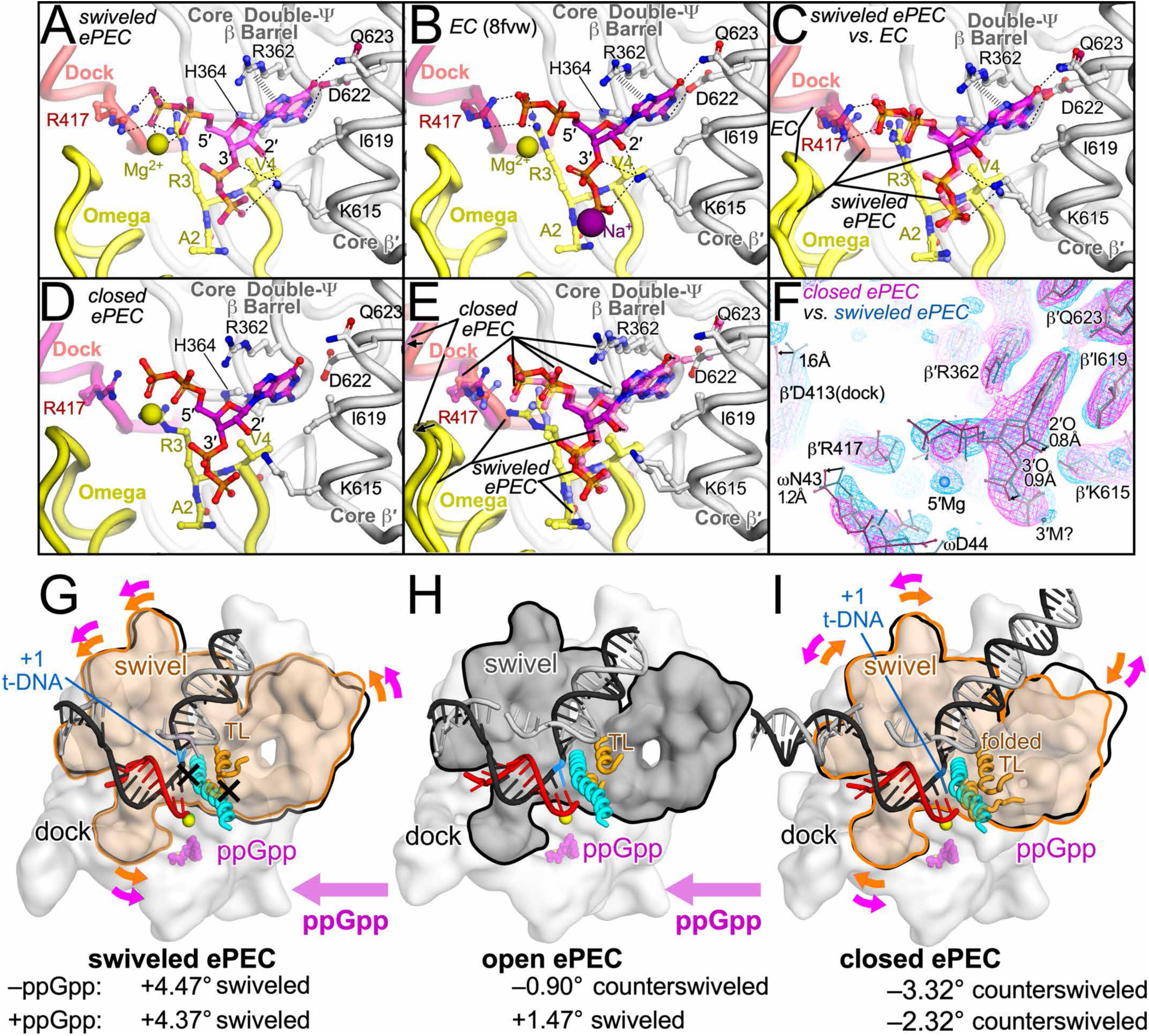
ppGpp binding to site 1 in swiveled versus counterswiveled ePECs and EC. (A) ppGpp bound to the swiveled ePEC (state *e* in Figure 2). Contacts are essentially identical to those reported for a stable EC (panel B).^27^ Key RNAP contacts to the ppGpp nucleobase are made by the immobile core of RNAP including the highly conserved double-λϑΙ β barrel whereas contacts to the phosphates are made by the dock (salmon) and ω (yellow) that change position during RNAP swiveling. Metal ions near ppGpp were weakly resolved in ePECs and only shown if assignment was possible. (B) ppGpp bound in the stable EC (PDB ID 8fvw) with metal ions as reported by Weaver et al., 2023.^27^ (C) Swiveled ePEC (panel A) and EC (panel B) structures aligned by the conserved double-λϑΙ β barrels in β and β′ subunits showing essentially identical ppGpp contacts. (D) ppGpp bound TL-folded, counter-swiveled ePEC (state *a* in Figure 2). (E) Swiveled (panel A) and counterswiveled (panel D) ePECs aligned as described for panel C. ppGpp is shifted away from the core RNAP contacts in the counterswiveled ePEC by ∼1.5 Å (indicated by black arrows on the 2′ and 3′ oxygens, 1.3 and 1.4 Å, respectively). Movements of the phosphates was accompanied by a shift in the Cαs of the dock and ω indicated by black arrows of 1.5 and 1.1 Å, respectively. (F) cryo-EM maps of the ppGpp-bound TL-folded ePEC (magenta) and swiveled ePEC (blue) aligned as in panel E and contoured to the same rmsd. The shifts in ppGpp, dock, and ω positions are evident in the aligned maps and denoted by dotted lines for ppGpp 2′ and 3′ oxygens, dock Asp413, and ωN43 (1.2, 1.4, 1.5, and 1.1 Å, respectively). The possible positions of metal ions are shown but are indistinguishable from water except as shown in panels A and D. (G) – (I) Comparison of swivel module (orange and gray) positions in the swiveled ePEC (G), open ePEC (H), and closed (counterswiveled) ePEC (I). Orange arrows indicate the rotation of the swivel module relative to its position in the open ePEC. Magenta arrows indicate the effects of ppGpp in favoring swiveling and disfavoring counterswiveling.

The differences in ppGpp contacts and effects on ePEC-state distributions are best understood in the context of the bi-directional model of RNAP swiveling when viewed into the RNAP active site (Figures 3G–3I; opposite to view of swivel module in Figure 2A). In this view, swiveling in the ePEC rotates the swivel module counter-clockwise, slightly kinks the BH, and inhibits TL folding by repositioning SI3.^20^ Inhibition of TL folding and of +1 t-DNA translocation by the kinked BH increases ePEC dwell time.^22^ Reinforcement of swiveling by ppGpp interaction with the dock and μ can explain ppGpp stimulation of pausing. The reduced ppGpp contacts in the counter-swiveled, closed ePEC may explain the ppGpp-induced shift away from TL-folded states. This inhibitory effect of ppGpp on counter-swiveling implies ppGpp might also affect nucleotide addition since counter-swiveling accompanies formation of the closed EC active state.^26,27^

### ppGpp alters RNAP occupancy and pausing in vivo

To understand how ppGpp acts on ECs and PECs in vivo, we next sought to distinguish direct effects of ppGpp on RNAP from indirect effects of ppGpp on translation and NTP levels. High ppGpp slows overall RNAP elongation rate in *E. coli* by a factor of ∼2,^33^ and even basal ppGpp may modestly stimulate RNAP pausing in vivo.^31^ However, direct versus indirect effects of ppGpp on cellular RNAP have not been deconvoluted. To enable quantitative comparison of elongating RNAP occupancies and pausing between strains and conditions with base-pair resolution, we developed quantitative NET-seq (qNET-seq) using a spike-in of *B. subtilis* cells expressing FLAG-tagged *Bsu*RNAP (Figure 4A and Table S2; Methods). We compared *E. coli* cells in which ppGpp was induced to mM level using an IPTG-inducible, constitutive RelA enzyme (RelA*) to cells with basal ppGpp levels in which expression of an inactive RelA variant (RelA^−^) was induced (Figure 4A).^34,39,48,49^

**Figure 4.**
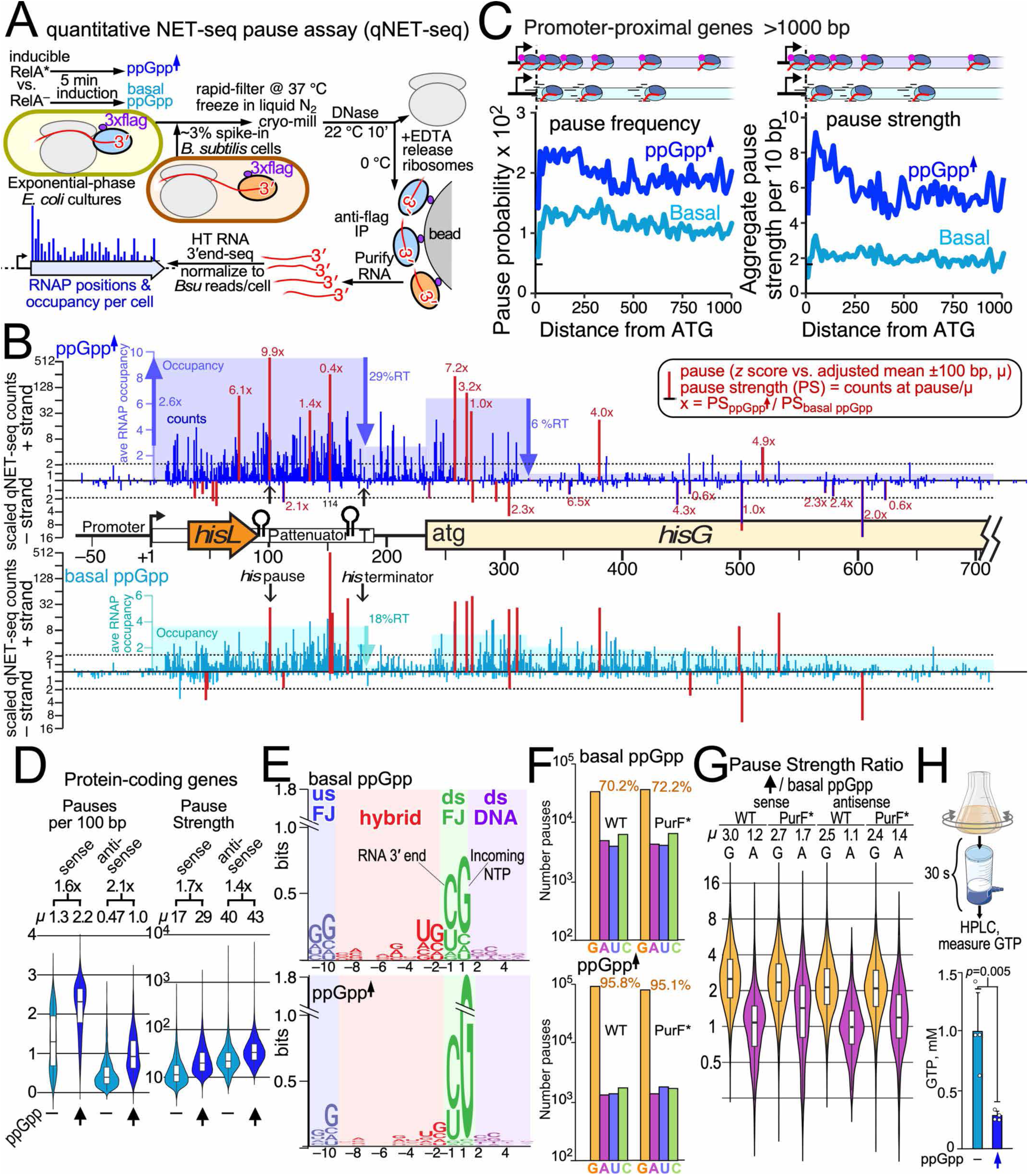
qNET-seq reveals that ppGpp primarily affects RNAP pausing before G in vivo. (A) Quantitative NET-seq (qNET-seq). *E. coli* cells with or without elevated ppGpp synthesis after IPTG induction of RelA variants that are constitutively active (RelA*) or inactive (RelA^−^) were harvested by filtration at 37 °C after an addition of a spike-in of ∼3% *B. subtilis* cells expressing 3xFLAG-tagged RNAP. Cells were collected, cryo-milled, and subjected to NET-seq analysis (Methods). *B. subtilis* nascent RNA reads were used to scale *E. coli* NET-seq signals to enable quantitative comparison among samples. (B) qNET-seq analysis of ppGpp effects on the *E. coli his* biosynthetic operon. High ppGpp (blue), basal ppGpp (teal). *hisL*, the leader peptide coding region. P and T indicate the positions of the *his* leader hairpin-stabilized pause and intrinsic terminator, respectively, encoded by the attenuator region. Sense and antisense scaled read counts are shown above and below the x-axis, respectively. Red lines and numbers indicate pause sites and the factor increase in pause strength caused by high ppGpp. Antisense pause calls used a lower count threshold to illustrate antisense *his* operon pauses where average antisense counts were much lower than sense counts. The light shaded areas indicate average RNAP occupancy (average 3′-end counts) in key regions. The blue up arrow and number indicate the effect of ppGpp on RNAP initiation whereas the blue and teal down arrows and numbers indicate locations of RNAP termination and the apparent % readthrough (%RT) of RNAP at the location. (C) Aggregate pause frequencies relative to translation start sites (shown as frequency per bp) and pause strengths (shown as aggregate strength in rolling 10 bp windows) for expressed promoter-proximal genes greater than 1 kb in length at basal and high ppGpp (teal and blue, respectively). Schematics depict RNAP occupancy at high versus basal ppGpp. (D) Pause frequency and pause strength distributions for expressed protein-coding genes >500 bp at basal and high ppGpp. Means for each distribution and fold effects of ppGpp are shown above the plots. (E) Sequence logos for all called pauses in WT *E. coli* with basal ppGpp (top) and induced ppGpp levels (bottom). (F) Numbers of pauses called before GTP (orange), ATP (pink), UTP (blue), and CTP (green) incorporation for WT and PurF* strains with basal ppGpp (top) and induced ppGpp levels (bottom). (G) Distributions of fold increase in pause strength caused by high ppGpp for pauses prior to GTP (orange) or ATP (pink) incorporation for coding regions and antisense, noncoding transcription for WT and PurF* strains. Mean fold increases (*µ*) are shown above plot. (H) GTP concentration measured in cell filtered for 30 s prior to organic solvent quench for PurF* strains with basal ppGpp (teal) and induced ppGpp levels (blue) (Methods).

The *his* biosynthetic operon illustrates the complex effects of ppGpp on RNAP occupancy and pausing (Figure 4B). The *his* operon is controlled by a ppGpp-stimulated promoter and a translation-coupled attenuator in its leader region.^50,51^ qNET-seq reads (vertical blue lines) reveal an ∼2.6-fold, ppGpp-induced increase in average RNAP occupancy in the initially transcribed attenuator region (+1 to +182) (transparent blue areas; Figure 4B). This increase is attributable to ppGpp-stimulated initiation.^36^ Occupancy drops at the attenuator termination point to ∼18% readthrough at basal ppGpp and ∼29% readthrough at high ppGpp. The ppGpp-induced increase in readthrough is attributable to ppGpp-induced stalling of ribosomes in *hisL*; efficient translation of *hisL* increases termination at the attenuator. Conversely, RNAP occupancy was elevated by ppGpp early in *hisG* and then dropped precipitously downstream of strong ppGpp-stimulated, AUG-proximal pauses to about ∼6% readthrough after ∼85 bp of *hisG*. At basal ppGpp, in contrast, RNAP paused less early in *hisG* and RNAP occupancy persisted at high level thereafter. A gradual decline in RNAP occupancy occurs on most *E. coli* genes during normal cell growth.^52^ The large drop in *hisG* occupancy at high ppGpp is attributable to ppGpp-inhibited translation leading to ppGpp-stimulated RNAP pausing, π-dependent termination, or both.^37,53^

Transcriptional pauses were identified and assigned pause strengths (PSs) by comparing nascent RNA 3′-end counts to an adjusted mean of counts in a +/–100 window (Figure 4B; Tables S3 and S4; Methods). ppGpp increased PS ∼10x at the hairpin-stabilized *his* leader pause (+102 nt) whereas other pauses were increased 3–7x (+159, 169, 381 nt), not affected (+273 nt), or even decreased in strength by ppGpp (+166 nt) (Figure 4B, Table S5) This range of effects on PS likely reflects different impacts of direct versus indirect ppGpp effects at distinct pause sites (i.e., via ppGpp binding to RNAP versus changes in translation or NTP levels).

ppGpp-enhanced pausing early in genes, where transcription–translation coupling is established, was clearly evident in aggregate profiles of promoter-proximal genes (Figure 4C, Table S7). Both aggregate pause frequency and pause strength increased in response to high ppGpp, giving broad peaks of pausing in the first ∼250 bp of these genes. A modest enrichment of pausing in this region also was evident at basal ppGpp. These data are consistent with increased AUG-proximal pausing when ppGpp slows translation initiation but could also reflect direct or other indirect effects of ppGpp.

To distinguish translation-mediated versus other effects of ppGpp, we examined antisense, non-coding transcription. Antisense transcription, which should not be affected by coupled translation, was readily apparent in the *his* operon (Figure 4B, Table S6). Although less prominent in the *his* regulatory region, antisense RNAPs were present at ∼21% and ∼17% of sense RNAPs in *hisG* (at basal and high ppGpp, respectively). This high level of antisense transcription was a general feature of *E. coli* protein-coding genes when assayed by qNET-seq (35 ± 21% and 27 ± 19% of sense RNAP in expressed genes at basal and high ppGpp) (Figure S5A). This number of antisense RNAPs in coding regions is greater than previously appreciated from total RNA measurements^54,55^ and was uncorrelated with sense RNAP transcription (Figure S5A), providing a convenient way to examine pausing uncoupled from translation. The frequency of pausing by antisense RNAPs was similar to sense RNAPs considering the reduced statistical power to detect pauses at the lower occupancies characteristic of antisense transcription (Figure S5B). High ppGpp increased both antisense pause frequency (2.1-fold) and pause strength (1.4-fold) similarly to sense RNAP (1.6-fold and 1.7-fold, respectively) (Fig. 4D, Table S8). We conclude that ppGpp affects antisense RNAP pausing in protein-coding genes similarly to sense RNAP pausing, indicating that ppGpp effects on translation are not the primary cause for increased RNAP pausing at high ppGpp.

### ppGpp affects RNAP pausing indirectly by reducing GTP concentration

Nearly all pauses detected at high ppGpp occurred before GTP addition (G pauses) (Table S3). Although pauses detected at basal ppGpp favored G over A pauses, similar to previous reports,^38–40,47^ this bias was dramatically enhanced at high ppGpp (Figure 4E). A ppGpp-induced reduction in GTP concentration combined with competitive inhibition by ppGpp binding in site 3 could explain this result.^9,34^ Therefore, we asked if the ppGpp-induced G-pause bias would decrease in a strain bearing a ppGpp-resistant variant of a key GTP biosynthetic enzyme, PurF (R45A; hereafter PurF*).^34^ PurF* ameliorates the decrease in GTP caused by high ppGpp.^34^ We catalogued pauses before G, A, U, and C at basal and high ppGpp in WT and PurF* strains. In both WT and PurF* strains, high ppGpp dramatically biased RNAP toward G pauses in protein-coding genes (to 95.8% and 95.1% G pauses for WT and PurF*, respectively; Figure 4F; Table S9). Since ppGpp affected G and A pauses similarly in vitro, we calculated ppGpp-induced enhancement of pause strength for G versus A pauses (Figure 4G; Table S9). ppGpp increased G pause strength ∼3.0 fold for WT and ∼2.7 fold for PurF* with a modest shift toward stronger A pausing in PurF*. A similar shift to lower G:A pause strength ratio was seen for antisense pausing, suggesting that PurF* modestly ameliorates but does not prevent the ppGpp-induced reduction in GTP:ATP ratio in cells.

To ask directly if GTP decreased upon ppGpp induction, we measured GTP in cells collected the same way as qNET-seq samples (by filtration at 37 °C). ppGpp induction decreased GTP from ∼1 mM to ∼275 µM (Figures 4H and S4C; Table S10). However, even at basal ppGpp, the filtered cells contained less GTP than reported for actively growing *E. coli* (2.7–4.9 mM).^56,57^ Further, GTP decreased more when filtration time was increased from 30 s to 2 min (to ∼140 µM and ∼110 µM for basal and high ppGpp PurF* cells, respectively). This effect of filtration time occurred with and without RelA* induction (Figure S5C, Table S10).

Low GTP is expected to increase G-pause strength, but it also could increase pausing indirectly by uncoupling ribosomes, since translation uses GTP. However, the similar effects of ppGpp on antisense pausing and the strong G-over-A bias (Figures 4D and 4G) indicate that ppGpp primarily affects pausing by reducing substrate GTP required for RNAP escape from G pauses. Further, our results suggest NET-seq preferentially captures RNAP at G pauses due to metabolic shifts that occur during cell filtration. Uncoupled translation may contribute to ppGpp-stimulated promoter-proximal pausing (Figure 4C), but uncoupling is not the primary reason ppGpp modulates most pauses.

### ppGpp slows RNAP allosterically at both pause and non-pause DNA positions

To test whether ppGpp primarily affects pausing via GTP levels or allosterically via binding to RNAP site 1 in vivo, we compared PurF* strains encoding wild-type RNAP (RNAP^+^) or an RNAP site 1^−^ mutant that abolishes ppGpp binding^36^ using both qNET-seq and quantitative, anti-RNAP ChIP-seq (qChIP-seq) (Figure 5A, Tables S11–S14). qNET-seq revealed a striking disconnect in the effects of RNAP site 1 on pausing versus RNAP occupancy. Loss of the allosteric ppGpp-binding site had little to no effect on in vivo measurements of transcriptional pausing but decreased the occupancy level of transcribing RNAP on genes. ppGpp induction increased G versus A pause bias and lowered GTP levels in the site-1 mutant similarly to an RNAP^+^ strain (Figures 5B and 5C; Tables S9 and S10). The site-1 mutant also exhibited similar pause frequencies and strengths to RNAP^+^, including when ppGpp induction increased RNAP pausing (Figure 5D, Table S15). In contrast, RNAP occupancy per gene calculated from qNET-seq decreased in the site 1^−^ strain relative to RNAP^+^ at both basal and high ppGpp (Figures 5E and S5, Tables S16–S18). qNET-seq revealed a modest increase in total RNAP occupancy on protein-coding genes associated with high ppGpp and a modest decrease in the corresponding antisense RNAPs (Figure S6). This high ppGpp-associated increase was not evident on longer genes (>1 kb, Figure 5E), consistent with the increase in pausing leading to increased π-dependent termination. This loss of RNAP on genes likely occurred during cell filtration for qNET-seq. qChIP-seq, which captures RNAP by rapid formaldehyde fixation before RNAP responses can occur,^58^ revealed that high ppGpp strongly increased occupancy on genes >1 kb prior to cell harvest (Figure 5F and Table S16; note that qChIP-seq cannot distinguish sense and antisense RNAPs). qChIP-seq also confirmed the reduced RNAP occupancy on genes overall in the site 1^−^ strain. An effect of cell harvest also was evident in the low occupancy of ribosomal RNA operons as measured by qNET-seq (Figure S6; Tables S18 and S19) since equal fractions

**Figure 5.**
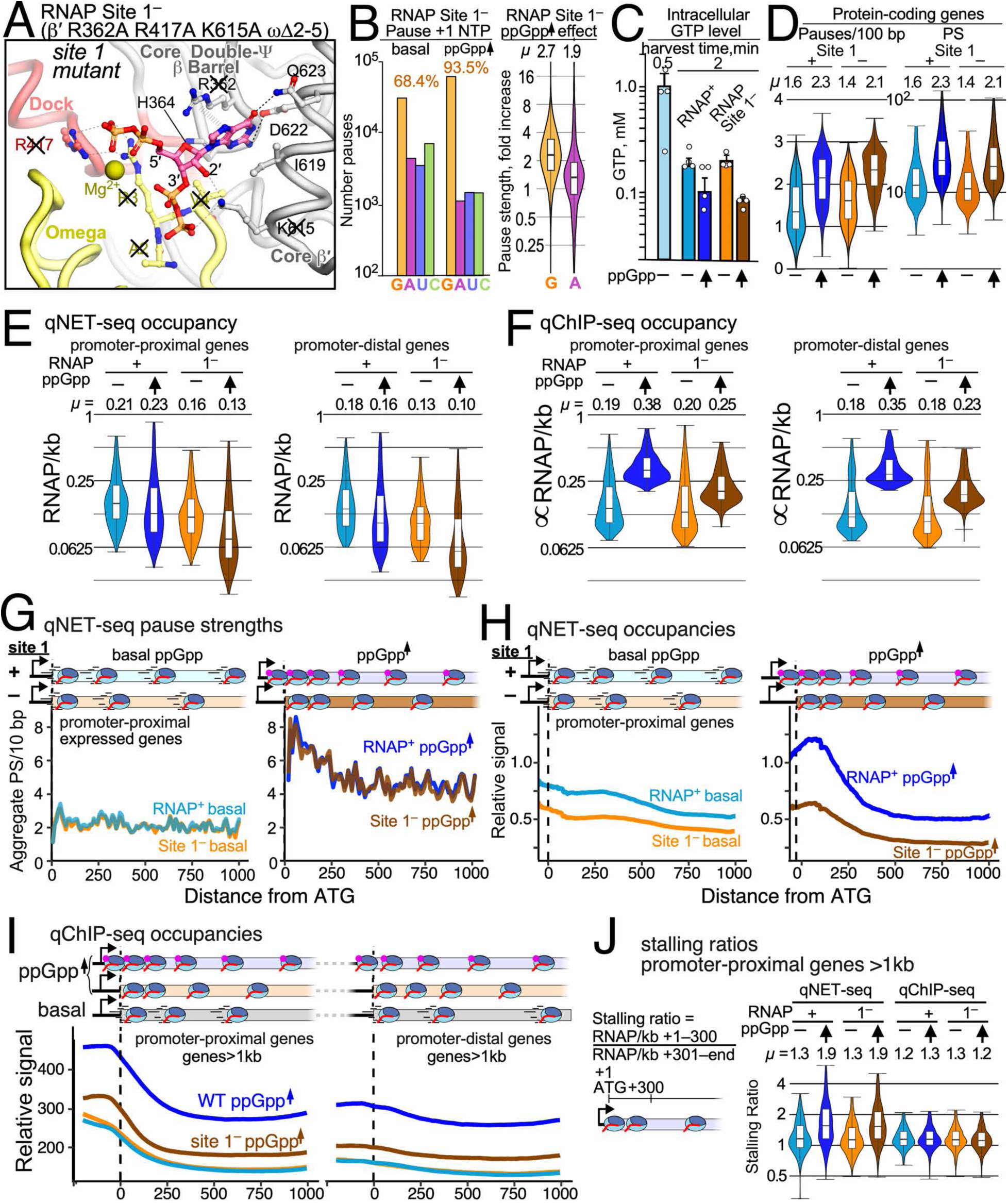
ppGpp allosterically slows RNAP broadly. (A) Amino-acid changes in the site 1^−^ mutant strain. Side chains marked with X are converted to Ala. (B) Number of pauses before G, A, C, or U incorporation for RNAP Site 1^−^ strain at basal and high ppGpp. Fold increase in strengths of pauses before G and A caused by high ppGpp in RNAP Site 1^−^strain. (C) Intracellular GTP levels measured for RNAP^+^ and Site 1^−^ strains with 2 min filtration at basal and high ppGpp. The level detected for RNAP^+^ at basal ppGpp with 0.5 min filtration (Figure 4H) is shown for comparison. RNAP^+^, basal ppGpp, teal; RNAP^+^, high ppGpp, blue; Site 1^−^, basal ppGpp, light orange; Site 1^−^, high ppGpp, dark orange. (D) ppGpp stimulation of pausing is largely unaffected by removal of ppGpp-binding site 1. The distribution of pause frequencies (left) and pause strengths (right) for expressed protein-coding genes >500 bp. Colors as in (C). (E) Distributions of qNET-seq RNAP/kb on promoter-proximal and promoter-distal genes >1 kb. Mean (*µ*) RNAP/kb is shown above the distributions. Colors as in (C). (F) Distributions of a qChIP-seq metric αRNAP/kb (proportional to RNAP/kb) on promoter-proximal and promoter-distal genes >1 kb. Mean (*µ*) RNAP/kb is shown above the distributions. Colors as in (C). (G) The qNET-seq distribution of pause strengths relative to translation start sites for expressed first genes in operons is unaffected by ppGpp. Basal ppGpp (RelA^−^ plasmid), left; high ppGpp (induced RelA* plasmid). RNAP^+^, teal and blue for basal ppGpp and high ppGpp levels, respectively; Site 1^−^ (dark orange and light orange for basal ppGpp and high ppGpp levels, respectively). (H) Relative occupancy of RNAP determined from qNET-seq signals for promoter-proximal genes greater than 1 kb. Basal ppGpp (RelA^−^ plasmid), left; high ppGpp (induced RelA* plasmid). RNAP^+^, teal and blue for basal ppGpp and high ppGpp levels, respectively; Site 1^−^ (dark orange and light orange for basal ppGpp and high ppGpp levels, respectively). (I) Relative occupancy of RNAP determined from qChIP-seq signals for promoter-proximal genes greater than 1 kb (left) and promoter-distal genes greater than 1 kb (right). RNAP^+^ and site 1^−^ at high ppGpp (induced RelA* plasmid; blue or light orange, respectively) or basal ppGpp (induced RelA^−^ plasmid, teal or dark orange, respectively). (J) The stalling ratio of RNAP on expressed promoter-proximal genes >1kb is not affected by removal of ppGpp-binding site 1 but is increased during cell harvest for qNET-seq analysis relative to cells rapidly fixed with formaldehyde for qChIP-seq analysis. Colors as in (C).

of RNAP are known to be present on protein-coding versus ribosomal RNA genes during cell growth in rich media.^59^

The discrepancy between no effect of ablating ppGpp–site 1 interaction on measures of RNAP pausing in vivo versus a decrease in RNAP occupancy in the site 1^−^ strain was also evident in aggregate plots of RNAP properties relative to translation start sites (Figures 5G–5I). High ppGpp increased pausing and accumulation of RNAP near start sites with no evident difference in pausing between RNAP^+^ and site 1^−^ (Figure 5G; Table S19). However, reduced RNAP occupancy in the site 1^−^ strain was evident throughout genes at both basal and high ppGpp (Figure 5H; Table S20). Reduced site-1^−^ RNAP occupancy also was evident by qChIP-seq when ppGpp was induced (Figure 5I, Table S20), excluding the possibility that it only reflected changes during cell harvest for qNET-seq. However, qChIP-seq indicated the effect of ablating site 1^−^ relied on high ppGpp since it was not evident at basal ppGpp (compare Figures 5H and 5I). An effect of qNET-seq cell harvest also was clearly evident in the buildup of RNAP near translation starts as measured by the stalling ratio of RNAP (fraction of RNAP in first 300 bp of promoter-proximal genes), which high ppGpp increased in qNET-seq but not qChIP-seq (Figure 5J, Table S21). We conclude that the strongest effect of high ppGpp on pausing arises from decreased substrate GTP rather than the allosteric effect of ppGpp, that loss of ppGpp binding to site 1^−^ decreases RNAP occupancy on genes, and that effects of ppGpp are greatly increased during cell harvest for qNET-seq.

## DISCUSSION

ppGpp control of transcript elongation by RNAP has long been known to be a crucial contributor to growth rate control and the stringent stress response in γ-proteobacteria. It also is a long-standing mechanistic mystery. By combining findings from classical biochemistry, cryo-EM, qNET-seq and qChIP-seq, we propose some answers to this mystery: (1) allosteric binding of ppGpp has two distinct effects on transcribing RNAP by biasing its swivel domain motions, inhibiting catalysis and prolonging pausing; (2) major ppGpp effects on pause escape rates in vivo are mediated by ppGpp-induced decreases in GTP levels; and (3) the allosteric effect of ppGpp on pausing in vivo may be masked by overall ppGpp effects on gene occupancy.

Swiveling was initially discovered in paused bacterial RNAPs as a counterclockwise rotation relative to ECs by ∼3.5–4.5° (as viewed from the β subunit side of RNAP) of about a third of the enzyme (the swivel module).^19,20,24,60^ Subsequently, smaller fluctuations of the swivel module in resting ECs and an ∼1.5° clockwise rotation (counter-swiveling) in the transition of ECs to the NTP-bound, folded-TL, RH–FL-down EC (closed EC) were reported.^26,30,61^ The closed EC closely resembles the pretranslocated, closed ePEC in which the folded TL contacts the 3′ RNA nucleotide instead of NTP (Figure 2).^22^ We suggest that these two different swiveling motions contribute to two distinct processes in RNAP (Figures 6A and 6B): (1) swiveling, which is an optional stabilization of offline paused states (e.g., as mediated by pause RNAP hairpins or NusA); and (2) counter-swiveling, which is an obligate motion in the transition between a relaxed EC and closed EC that positions NTP for catalysis during the nucleotide addition cycle (NAC).

**Figure 6.**
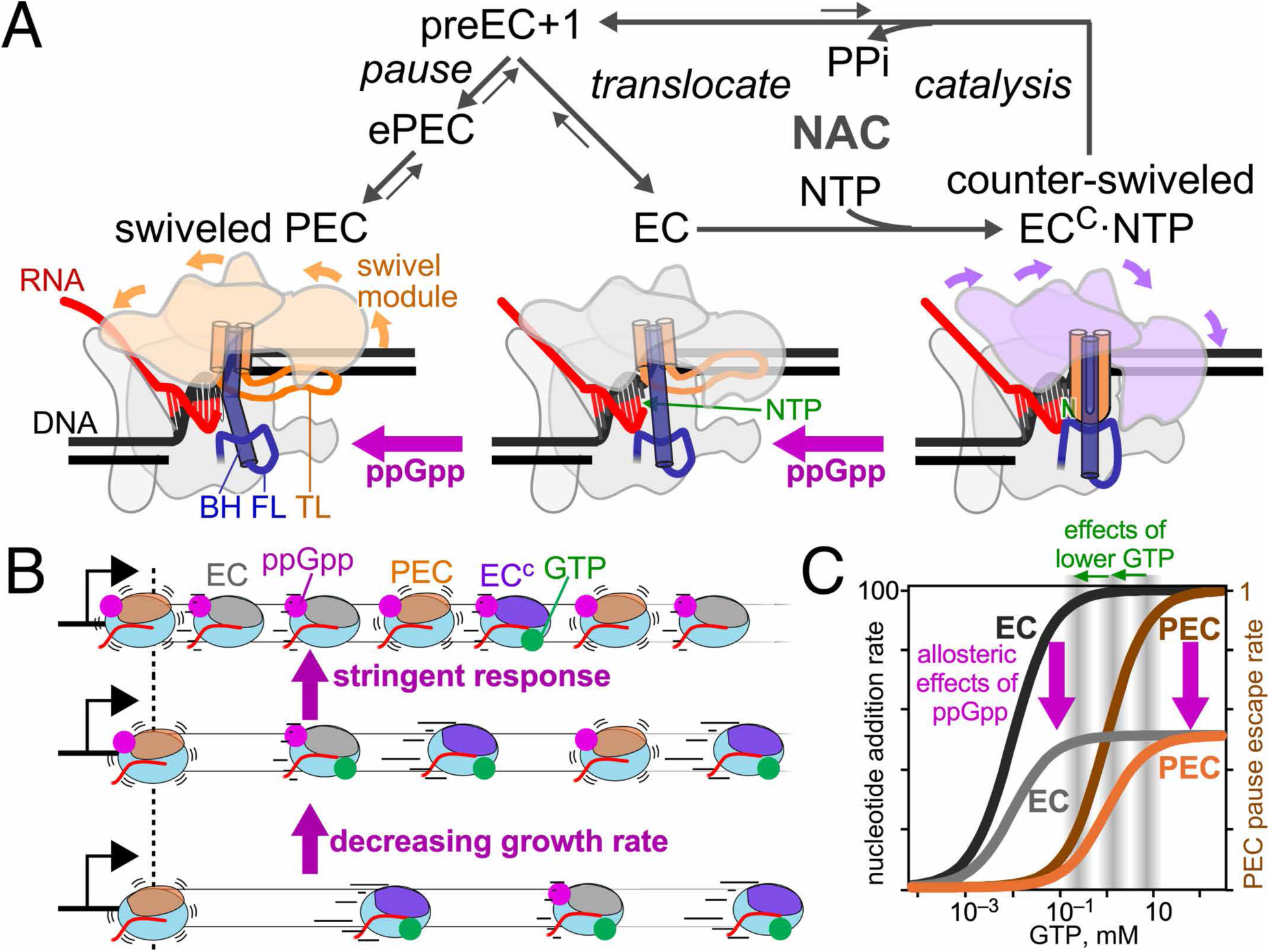
Model of ppGpp effects on two modes of RNAP swiveling. (A) The nucleotide addition cycle (NAC) including transcriptional pausing showing proposed effects of ppGpp on swiveled PEC and counter-swiveled closed EC (EC^C^). Movements of the swivel module in two different directions relative to the EC are shown by colored arrows (orange, toward swiveling; lavender, toward counter-swiveling). (B) Conceptual model of how changes in ppGpp during normal cell growth at different rates (decreasing growth rate regulation and in the stringent response (e.g., acute amino acid starvation) alter the populations of EC–PEC intermediates and occupancy of RNAP on genes. (C) Conceptual illustration of the different effects of changes in substrate GTP concentration and allosteric effects of ppGpp on pausing and the NAC. Idealized rates are shown without and with ppGpp for EC nucleotide addition (black and gray, respectively) and PEC pause escape (light and dark orange, respectively). The gray zones illustrate how lowering GTP concentrations can slow PEC escape with less effect on EC nucleotide addition, leading to increases in pause strengths measured by qNET-seq. The magenta arrows illustrate how the allosteric effect of RNAP on the NAC and on pausing via different modes of swiveling can increase RNAP occupancy without large effects on pause strengths measured by qNET-seq.

ppGpp affects both swiveling and counter-swiveling, as evidenced by ppGpp promotion of ePEC swiveling and inhibition of TL folding (Figures 1 and 2). Cryo-EM provides the strongest evidence for the two separate ppGpp effects. ppGpp quantifiably shifts ePEC conformations away from TL folded states and toward the fully swiveled ePECs (Figures 2D and 2E). Our CTR disulfide crosslinking results are consistent with distinct ppGpp effects on the transition from the open, unswiveled EC to the closed, counter-swiveled EC and on the transition from the unswiveled PEC to the swiveled PEC but cannot separate the two effects because the crosslinks report only the closed, counter-swiveled EC and swiveled PEC states.

This two-facet model for allosteric ppGpp effects on RNAP swiveling is attractive because it can explain an otherwise confusing relationship of swiveling to nucleotide addition and pausing. Swiveling has been proposed to be an obligate step in the NAC required either for translocation or for TL folding,^60,61^ and, alternatively, to occur in some but not all PECs that themselves form at a small fraction of DNA template positions.^19,20,22,24^ Assigning two distinct swivel motions, one obligate involved in the EC-to-closed EC transition and one optional involved in transcriptional pausing, explains how swiveling can be both obligate in every round of nucleotide addition and an optional aspect of pausing without contradiction. They are two different motions of RNAP that involve the same module.

In cells, ppGpp affects multiple processes (Figure 1A). Allosteric modulation of RNAP appears to be limited to certain bacterial lineages like γ-proteobacteria. In these lineages, ppGpp allosterically affects transcription initiation principally through site 2 formed by binding of DksA to RNAP and elongation principally through site 1 studied here.^12,27,31^ Because ppGpp also affects GTP and ATP levels,^62^ which modulate transcription initiation, and competes for GTP in the RNAP active site (ppGpp site 3), the ppGpp-mediated stringent response may rely in significant part on the indirect decrease in GTP levels. We confirm that this indeed is the case for the effect of ppGpp on transcript elongation by RNAP. In vivo, pausing before G is greatly strengthened when ppGpp is induced and GTP levels drop, whereas pausing before A is not strengthened to the same extent, unlike in vitro (Figures 2B and 4D-F). Reduced GTP can increase pausing with lesser effects on elongation due to differences in their relative *K*_GTP_, whereas an allosteric effect on swiveling may affect both pausing and elongation (Figure 6C).

The two-facet model for ppGpp effects on RNAP swiveling may also explain why we see little effect of RNAP site 1 on pausing in vivo as measured by qNET-seq (Figures 6B and 6C). Both qNET-seq and qChIP-seq revealed a site 1–dependent increase in transcribing RNAP occupancy on genes at high ppGpp (Figures 5E, 5F, 5H, and 5I). RNAP occupancy on genes inversely correlates with elongation rate when initiation is unchanged.^63^ Since pause frequency and strength calculations rely on signal changes relative to the mean of nearby NET-seq signal, slower elongation caused by ppGpp inhibition of the NAC may increase the mean, masking the effect on pause signals (Figure 6B). This relatively uniform effect of ppGpp differs from effects mediated by reducing GTP concentration because non-pause sites may exhibit little change due to a much tighter *K*_NTP_ than for pause sites (Figure 6C).^39,64^ Removal of site 1, unlike changes in GTP level, may give no net effect on pause frequency or strength because both the surrounding mean and pause NET-seq signals decrease. Thus, the reduced gene occupancy caused by removal of site 1 without extensive changes in measured pause strengths is consistent with the two-facet model for ppGpp effects on RNAP swiveling. However, we cannot exclude the possibility that the site 1^−^ mutant instead somehow increases dissociation of RNAP during transcription when ppGpp is elevated.

Although ppGpp-stimulation of backtracking does not explain ppGpp-stimulated transcriptional pausing (Figure 1D and S1), the two-facet model of ppGpp–RNAP allosteric action remains compatible with reported effects of ppGpp on transcription-coupled DNA repair (TCR) and the related phenomenon of stress-induced DNA mutagenesis.^8,65,66^ ppGpp-induced backtracking is proposed to aid both TCR and mutagenesis. Swiveling is associated with backtracking; thus, ppGpp may increase backtracking via effects on swiveling.^26^ However, an increased propensity for backtracking can occur without increasing pause dwell times because rapid changes in RNA register may not be rate limiting for pause escape, as found for the consensus ePEC.^21^

### Limitations of study

Our results provide important new insights into ppGpp direct (allosteric) and indirect effects on transcription and plausible explanations for how ppGpp may affect two facets of RNAP swiveling. However, our detection of the ppGpp effect on TL folding relies on distributions of ePEC conformations that include TL-folded states and not direct detection of ppGpp action on the NAC. Additionally, computational sorting of particles to distinct conformational states during cryo-EM data processing is inherently limited by the signal-to-noise ratio of the underlying individual particle images. Our assignment of direct effects of ppGpp on pausing in vivo relies on statistical analyses using spike-in samples. Although qNET-seq and qChIP-seq are improvements over existing methods commonly used for bacterial studies, they cannot fully control for pleiotropic effects of ppGpp and changes in ECs and PEC distributions that occur during sample workup. Our measurements of GTP levels revealed that GTP decreases during NET-seq cell harvest (Figures 4H, S4, and 5C), highlighting the importance of rapid RNAP inactivation for accurate detection of RNAP positions.^47^ Direct genome-scale measurements of transcription in vitro could provide more controlled conditions for these types of mechanistic analyses.

## Resource Availability Lead Contact

Further information and requests for resources and reagents should be directed to and will be fulfilled by the Lead Contact, Robert Landick.

## Materials Availability

All unique reagents generated in this study are available without restriction from the Lead Contact.

## Data and Code Availability

- Model and map files were deposited at the Protein Data Bank (PDB) and Electron Microscopy Data Bank (EMDB), respectively: ePECswiveled+G4P (PDB 12EB, EMDB EMD-76371), ePECopen+G4P (PDB 12ED, EMDB EMD-76373), ePECsemi-closed+G4P (PDB 12EF, EMDB EMD-76375), ePECclosed+G4P (PDB 12EH, EMDB EMD-76377), ePECswiveled (PDB 12EC, EMDB EMD-76372), ePECopen (PDB 12EE, EMDB EMD-76374), ePECsemi-closed (PDB 12EG, EMDB EMD-76376), ePECclosed (PDB 12EI, EMDB EMD-76378). They are publicly available as of the date of publication.
- All DNA sequencing data and primary mapped reads have been deposited to NCBI GEO GSE327058 and are publicly available as of the date of publication.
- All original code has been deposited on Zenodo and is publicly available at doi: 10.5281/zenodo.19392801 as of the date of publication.
- Any additional information required to reanalyze the data reported in this paper is available from the Lead Contact upon request.

## Acknowledgements

The authors are grateful for extensive discussions during the course of this work with colleagues in the Landick, Darst, and Elizabeth Campbell labs. We thank Wilma Ross and Rick Gourse for sharing strains and plasmids essential to this study and David Stevenson and Daniel Amador-Noguez for help analyzing cellular GTP levels. This work was supported by NIH grants R01 GM038660 to R.L. and R35 GM118130 to S.A.D. Cryo-EM data reported here were collected at the Simons EM Center and the National Resource for Automated Molecular Microscopy located at the New York Structural Biology Center, supported by grants from the National Institutes of Health (NIH) National Institute of General Medical Sciences (P41 GM103310), NYSTAR, the Simons Foundation (SF349247), the NIH Common Fund Transformative High-Resolution Cryo-Electron Microscopy program (U24 GM129539) and the NY State Assembly Majority.

## Author contributions

Conceptualization, R.L. and S.A.D.; Methodology, A.M, R.M., Y.B., S.D., M.E., and R.L.;

Investigation, A.M., R.M., M.E., Y.B., J.B., J.S., and J.L.; Data Curation, A.M., M.E., M.W.,

B.S., and R.L.; Formal Analysis, A.M., M.W., and R.L.; Software, M.W., M.E., and R.L.; Visualization, R.L. and A.M.; Writing – Original Draft, R.L.; Writing – Review & Editing, R.L., A.M., R.M., M.E., and S.A.D.; Funding Acquisition, R.L. S.A.D.

## Declaration of interests

The authors declare no competing interests.

## Supplemental Information Index

Document S1. Contains Figures S1-S5.

## METHODS

## Key Resources Table

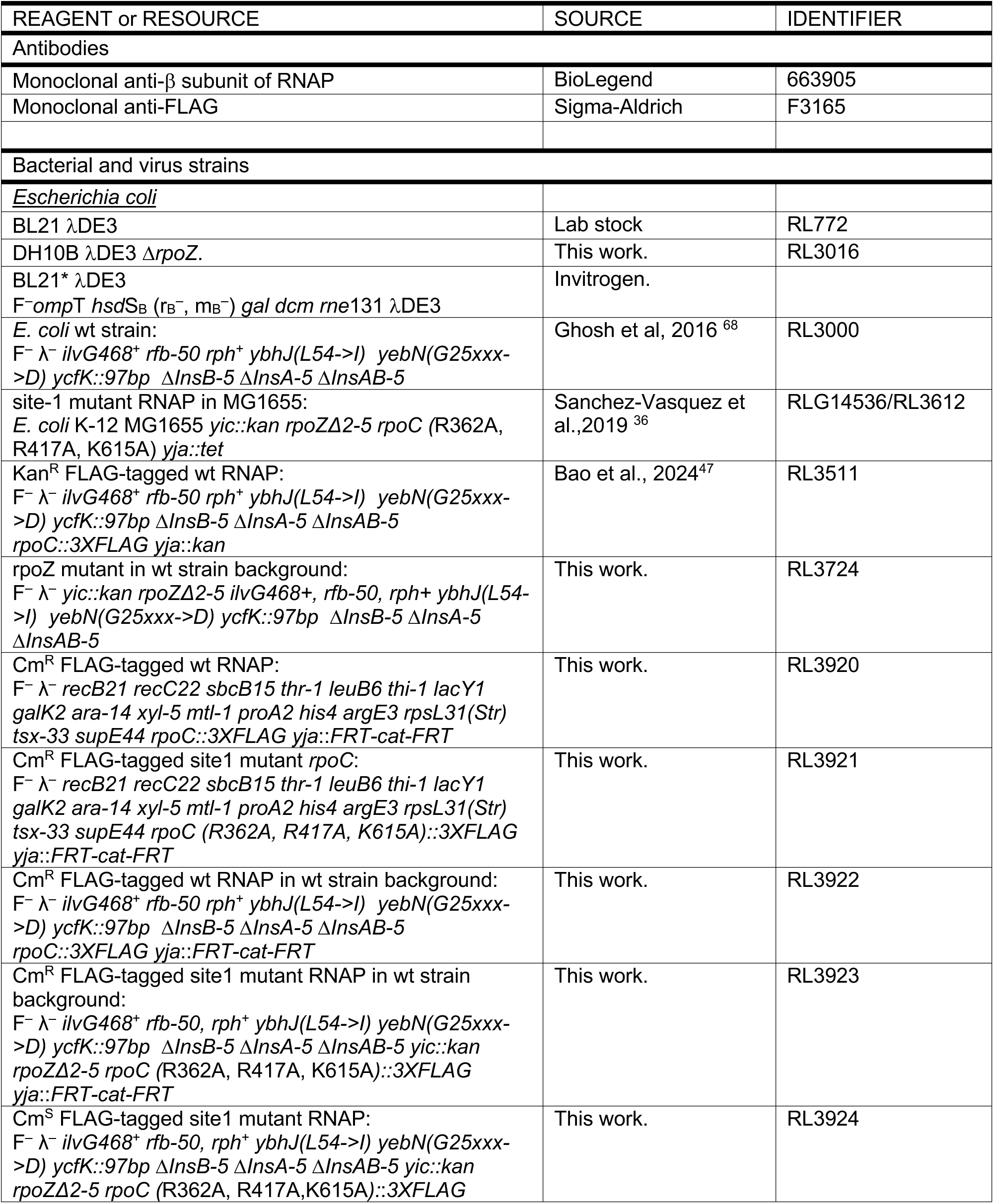
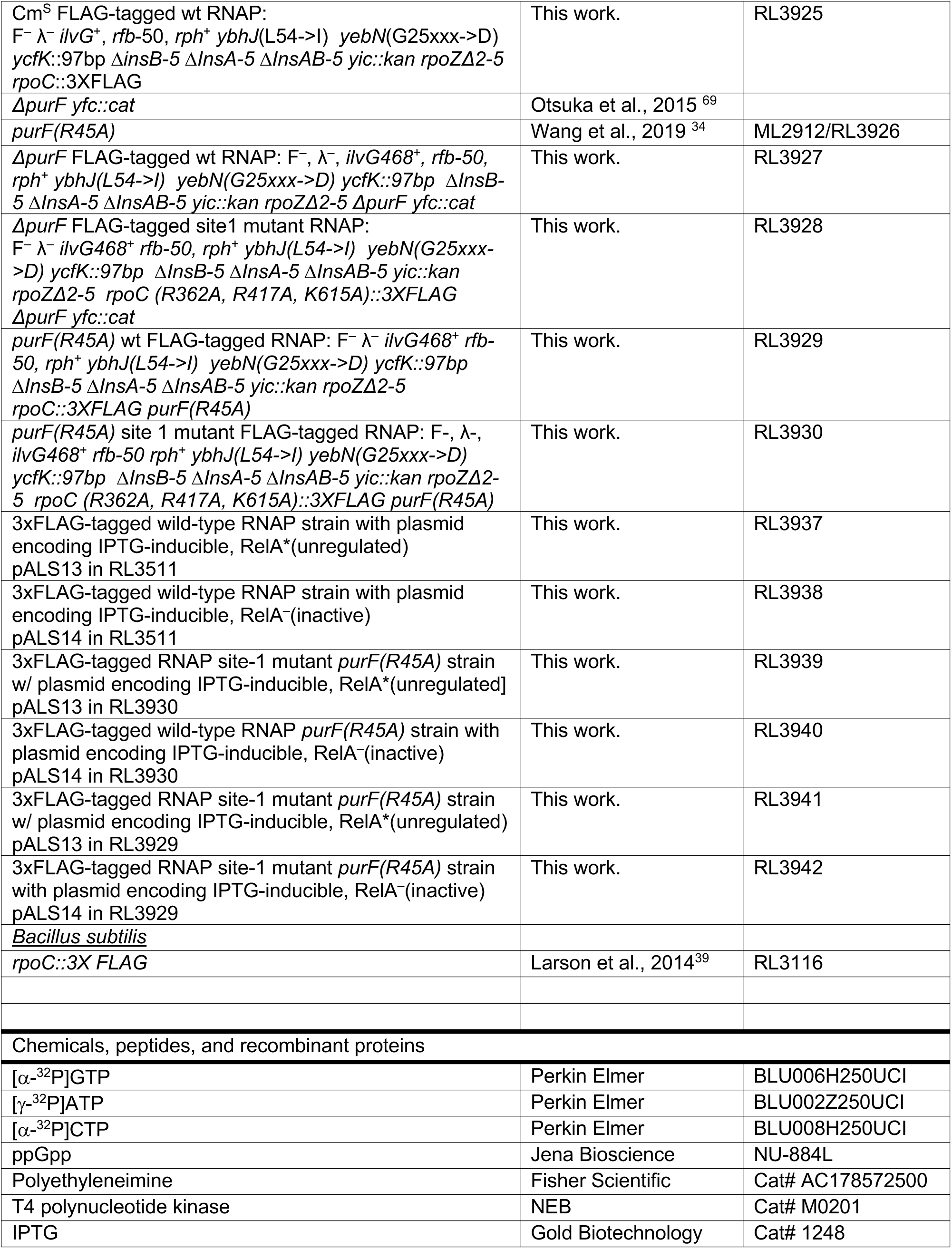
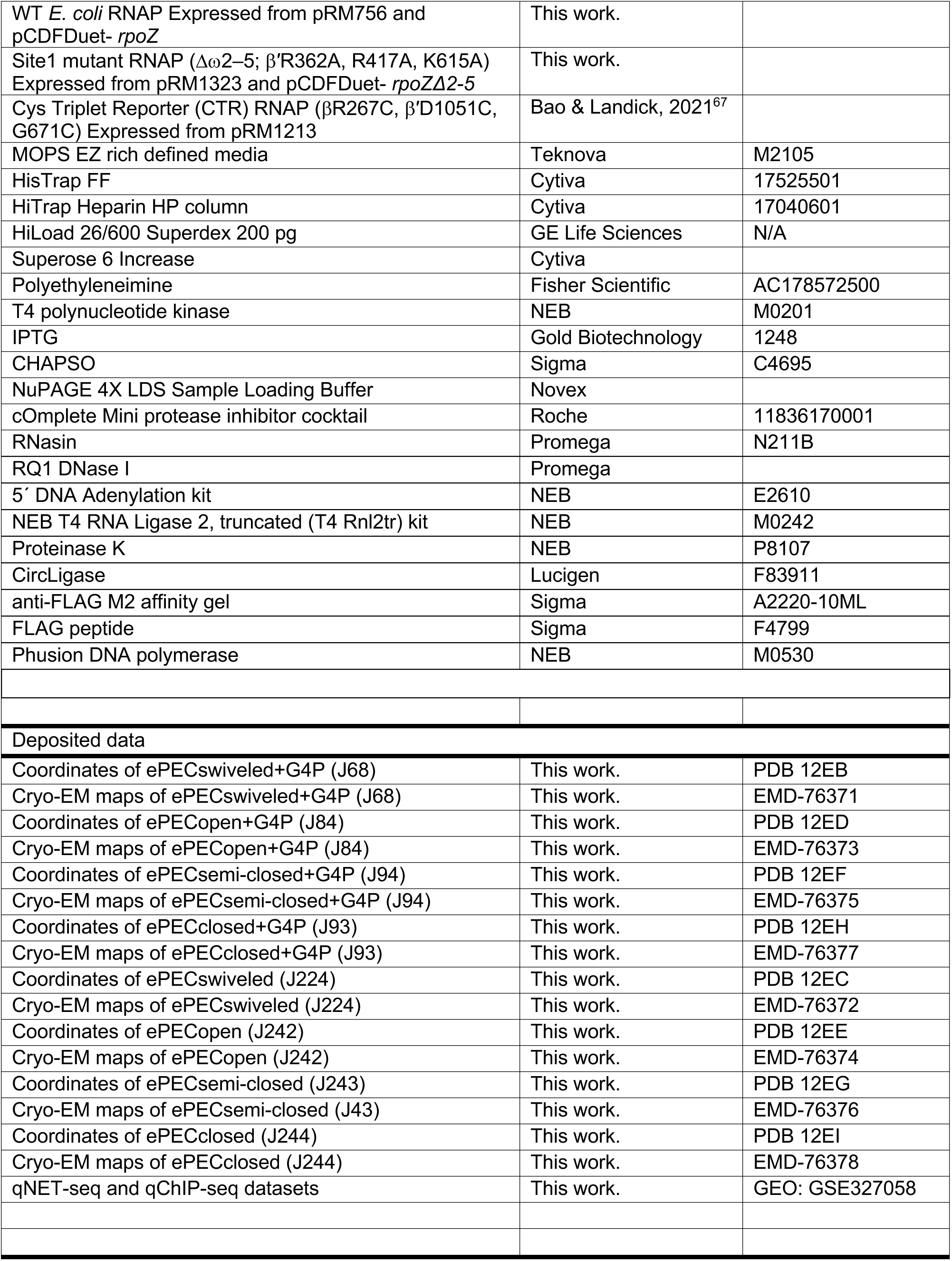
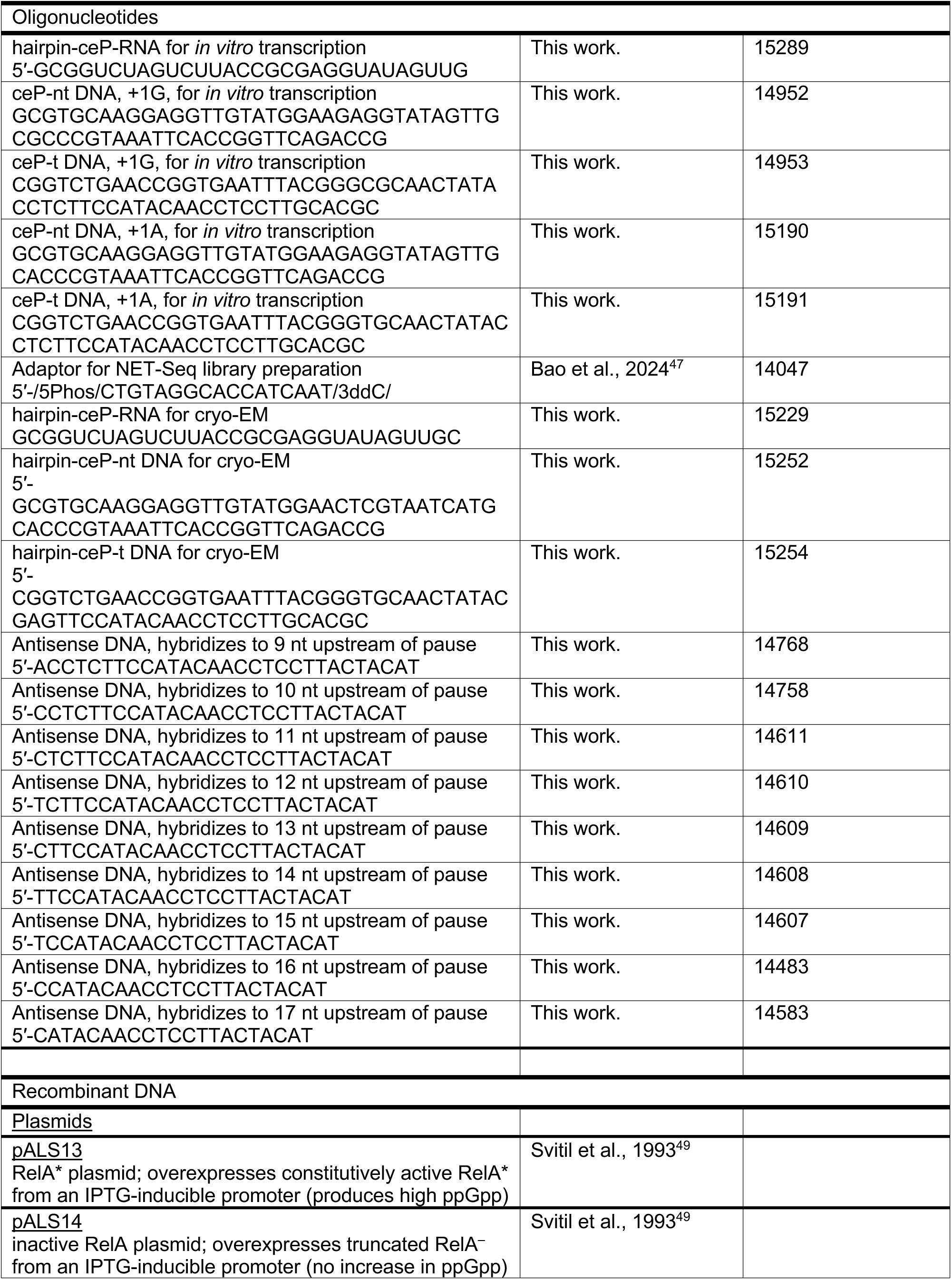
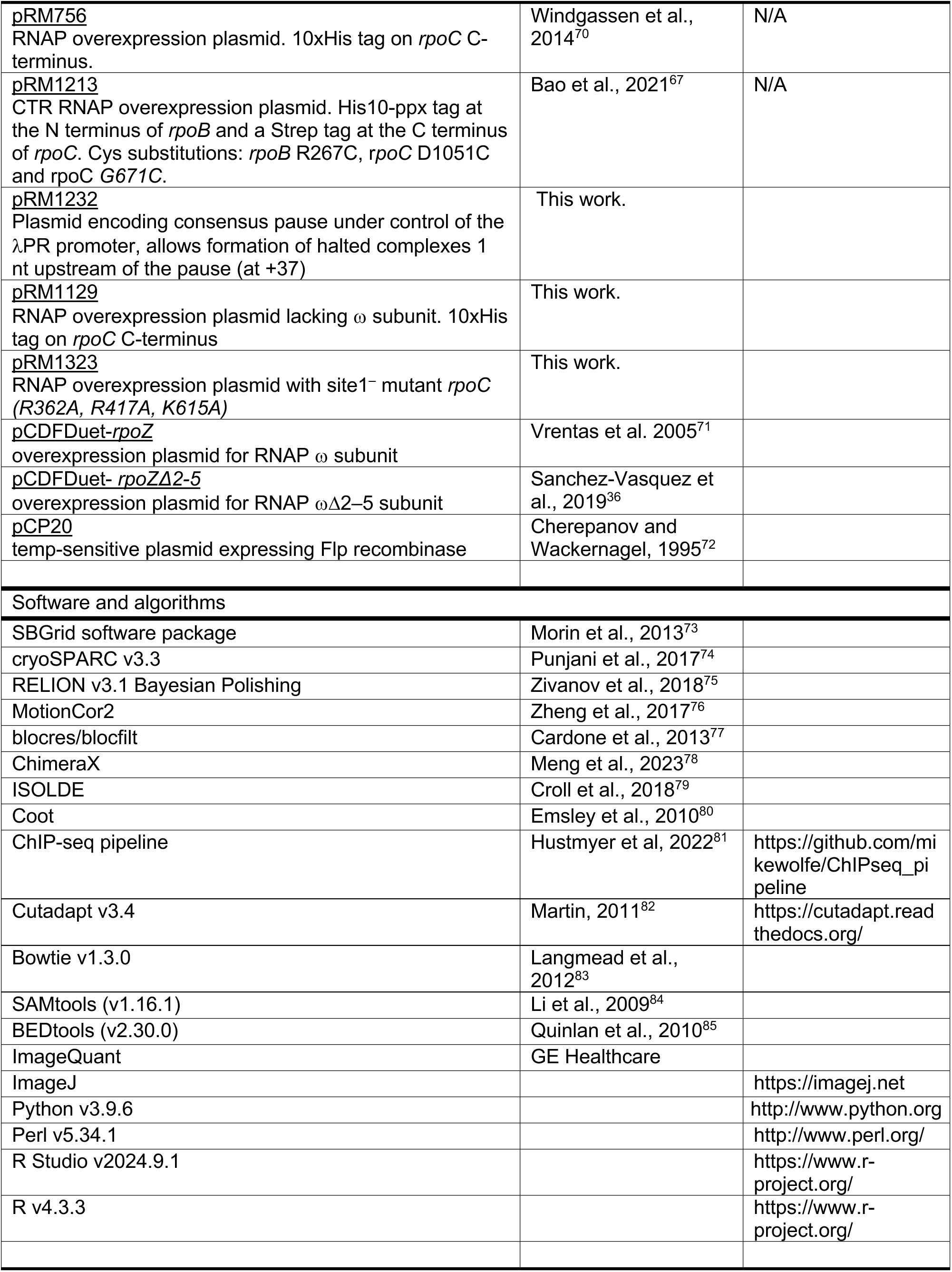

## EXPERIMENTAL MODEL AND STUDY PARTICIPANT DETAILS

### Bacterial strains and cultivation

*E. coli* strains containing FLAG-tagged wild-type or site-1^−^ RNAP were constructed in several steps. First, plasmids containing either 3xFLAG-tagged wildtype *rpoC* or FLAG-tagged site-1 mutant *rpoC* (R362A, R417A, K615A) were used as recombination templates to produce strains RL3920 and RL3921 with 3xFLAG-tagged *rpoC* and nearby chloramphenicol (Cm) resistance markers (*FRT-cat-FRT*). Next, the 3xFLAG-tagged *rpoC* genes were moved by P1 transduction into the wild-type *E. coli* strain RL3000 or RL3000 containing an N-terminal deletion of *rpoZ* (RL3724, *yic::kan rpoZΔ2-5*; from RLG14536/RL3612)^36^ by selection for Cm resistance to give strains RL3922 and RL3923, respectively. The Cm-resistance markers were removed using FLP recombinase using temperature-sensitive plasmid pCP20^72^ to give strains RL3924 and RL3925. To introduce the *purF* variant (*purF* R45A), a deletion of *purF*^69^ was first introduced to these strains by P1 transduction and selection for Cm resistance, giving auxotrophic strains RL3927 and RL3928. These strains were then transduced with P1 grown on the *purF* R45A strain by selecting for transductants able to grow on minimal media, resulting in strains RL3929 and RL3930.

To control the levels of ppGpp, plasmids encoding IPTG-inducible, constitutively active (RelA*) or inactive (RelA^−^) forms of the ppGpp synthetase RelA (pALS3 and pALS14, respectively) were transformed into RL3511 (strains RL3937 and RL3938) or RL3929 and RL3930 (strains RL3939-RL3942) with selection for ampicillin.

### Cell culture

For NET-Seq and ChIP experiments, a single colony was inoculated into MOPS EZ rich defined medium with 0.2% (w/v) glucose (RDM, Teknova) for overnight growth at 37 °C. A portion of the overnight culture was inoculated to apparent 0.05 OD_600_ into 0.85 L RDM supplemented with ampicillin (100 µg/mL) to maintain plasmids pALS13 or pALS14. The cultures were grown in Fernbach flasks on an orbital shaker at 200 rpm and 37 °C until cell density reached early log-phase (apparent OD_600_ ∼0.39 +/– 0.02). Expression of unregulated RelA**\*** or inactive RelA^−^ was induced by addition of IPTG to 1 mM with continued shaking at 37 °C. After 5 minutes of induction, cells for GTP quantitation were removed and rapidly filtered (described in detail below). Next, 50 mL samples were removed for ChIP, and the remaining culture for NET-Seq analysis was mixed with FLAG-tagged RNAP *B. subtilis* cells (strain RL3116) as spike-in and rapidly filtered through a 0.45 µm cellulose filter. The collected cells were quickly scraped from the filter with a spatula, and plunged into liquid nitrogen before storage at –80 °C as described previously.^48^ Cells for ChIP were immediately mixed with sodium phosphate (pH 7.6, 10 mM final) and formaldehyde (1% final) and incubated at 37 °C for crosslinking as described previously,^86^ followed by quenching with glycine (100 mM final) for 30 min at 4 °C. The crosslinked cells were then centrifuged, washed multiple times with phosphate-buffered saline (PBS; 137 mM NaCl, 2.7 KCl, 10 mM Na_2_HPO_4_, 1.8 mM KH_2_PO_4_), and stored at –80 °C. Biological duplicate samples were collected for each strain or experimental condition.

*B. subtilis* cells were added from pre-measured and stored aliquots to *E. coli* cultures immediately before harvest at a ratio of 3.3% *B. subtilis*-to-*E. coli* cells as calculated from a pre-determined relationship between apparent OD_600_ and colony-forming units for each *E. coli* strain grown in RDM. Aliquots of *B. subtilis* RL3116 cells at measured cell densities were prepared from late log–phase cultures that were cooled on ice, mixed with cold glycerol to a final concentration of 15% glycerol, flash-frozen in liquid nitrogen, and stored at –80°C. Prior to adding *B. subtilis* cells to the *E. coli* cultures, aliquots were thawed in ice water and stored on ice.

## METHODS DETAILS

### Purified RNAPs

RNAPs for transcription assays and CTR assays were purified as described previously ^47^. Briefly, Wildtype (WT) *E. coli* RNAP was expressed from pRM756 and pCDFDuet- rpoZ in BL21 λDE3 or BL21* λDE3. Site1^-^ RNAP (Δρο2–5; β′R362A, R417A, K615A) was expressed from pRM1323 and pCDFDuet- rpoZΔ2-5. Cys Triplet Reporter (CTR) RNAP (β R267C, β ′D1051C, β G671C) was expressed from pRM1213. After transformation of RNAP expression plasmids, a single colony was inoculated into 5 mL LB with 50 µg kanamycin/mL or 5 mL LB with 50 µg kanamycin/mL and 50 µg spectinomycin/mL and grown overnight at 37 °C. The saturated cell culture was then added to 1-2 L fresh LB supplemented with the same antibiotics and grown at 37 °C with adequate aeration by orbital shaking in a Fernbach flask until apparent OD_600_ reached 0.6. Protein expression was induced by adding IPTG (Gold Biotechnology) to 1 mM and cell growth was continued for 3 hours at 37 °C or overnight at 16 °C. The cells were harvested and homogenized by sonication in 30 mL Lysis Buffer (50 mM Tris-HCl pH 7.9, 5% v/v glycerol, 233 mM NaCl, 2 mM EDTA, 10 mM β-mercaptoethanol, 10 mM DTT, 100 µg/mL PMSF, and 1 tablet of protease inhibitor cocktail (Roche cOmplete, Mini, EDTA-free). After removing cell debris by centrifugation (11000 × *g*, 15 min, 4 °C), DNA binding proteins including target RNAPs were precipitated by addition of polyethylenimine (PEI, Sigma-Aldrich) to 0.6% (w/v) final. After centrifugation (11000 × *g*, 15 min, 4 °C), the protein pellet was resuspended in 25 mL PEI Wash Buffer (10 mM Tris-HCl pH 7.9, 5% v/v glycerol, 0.1 mM EDTA, 5 µM ZnCl_2_, 500 mM NaCl) to remove non-target proteins. After centrifugation (11000 × *g*, 15 min, 4 °C), RNAP was eluted from the pellet into 25 mL PEI Elution Buffer (10 mM Tris-HCl pH 7.9, 5% v/v glycerol, 0.1 mM EDTA, 5 µM ZnCl_2_, 1 M NaCl). The crude extract of RNAP was subjected to sequential FPLC purifications using Ni^2+^-affinity (HisTrap FF 5 ml, GE Healthcare Life Sciences) and heparin-affinity (Heparin FF 5 ml, GE Healthcare Life Sciences). The purified RNAP was dialyzed into RNAP Storage Buffer (10 mM Tris-HCl, 25% v/v glycerol, 100 mM NaCl, 100 µM EDTA, 1 mM MgCl_2_, 20 µM ZnCl_2_, 1-10 mM DTT) to final concentrations of 5–10 mg/mL. The higher concentration of DTT (10 mM) was included in the CTR RNAP storage buffer to inhibit disulfide bond formation by engineered cysteine residues.

### *In vitro* transcription assays

In vitro transcription assays to measure pausing were performed as described previously with minor modifications.^47^ For scaffold-based assays of the consensus elemental pause (Figure 1), pre-assembled nucleic-acid scaffolds of RNA and template strand (t-DNA) were incubated with *E.coli* RNAP core enzyme in Transcription Buffer (10 mM Hepes, pH 7.9, 50 mM potassium glutamate, 10 mM magnesium glutamate, 0.1 mM EDTA, 1 mM DTT and 5 µg acetylated BSA/mL) at final concentrations of 0.5 µM RNA, 1.0 µM t-DNA, and 1.5 µM RNAP for 15 min at 37 °C. Non-template strand (nt-DNA) was added to 1.5 µM and the mixture was incubated for an additional 15 min at 37 °C. (Sequences of oligos used to assemble the ePEC with incoming GTP or ATP are listed in the Resource Table.) Further reconstitution was then blocked by incubation with heparin (0.1 mg/mL final) for 5 min at 37 °C. To label the complexes, 150 nM reconstituted ECs were labeled with [α-^32^P]CMP and halted one nucleotide before the pause (–1). These complexes were then reacted with 200 µM GTP or ATP, with or without ppGpp (100 µM). Samples were removed at different time points and mixed with an equal volume of Stop Buffer (8 M urea, 50 mM EDTA, 90 mM Tris-borate buffer, pH 8.3, 0.02% bromophenol blue, and 0.02% xylene cyanol). RNAs were resolved by denaturing PAGE (8% polyacrylamide, 19:1 acrylamide:bis-acrylamide, 45 mM Tris-borate, pH 8.3, 1.25 mM Na_2_EDTA, 8 M urea), exposed to a Storage Phosphor Screen (GE Healthcare) and scanned with Typhoon PhosphorImager (GE Healthcare). An ^32^P-labeled pBR322 MspI digested DNA ladder (NEB) was run on the gel for size reference.

To assess a potential role of ppGpp in the suppression of backtracking at the elemental consensus pause, *in vitro* transcription experiments were performed using a promoter-initiated template (pRM1232) allowing formation of halted complexes one nucleotide before the pause (G36 ECs). Halted complexes were generated by mixing 50 nM RNAP, 25 nM double-stranded template, 125 µM ApU dinucleotide, and 2.5 µM ATP, UTP, and [α-^32^P]GTP and incubating the reaction for 10 min at 37 °C. To test effects of backtracking, different antisense DNA oligonucleotides (asDNAs) were paired to the nascent RNA. These asDNAs were added to the halted G36 ECs with or without ppGpp (100 µM) and allowed to incubate for 3 min at 37 °C before addition of 100 µM all four NTPs to allow elongation through the pause. As described above, samples were removed at different time points and mixed with an equal volume of Stop Buffer and separated by denaturing PAGE.

All gels were quantified using ImageQuant software (GE Healthcare).

To analyze pause kinetics, the pause fraction change through time was fit with bi-exponential decay equation:

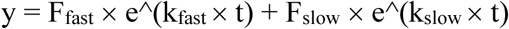

F_fast_ and F_slow_ represent the fraction of RNA species entering the slow and fast escape phase and *k*_fast_ and *k*_slow_ describe their escape rates. The pause strength (τ) was calculated as:

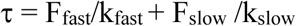

### Cys-triplet CTR conformation assay, gel quantitation, and SPB calculation

The Cys-triplet (CTR) assay was used to measure the extent of swiveling in ePECs with and without bound ppGpp (100 µM). The DNA–RNA partially noncomplementary scaffold was assembled by incubation of purified oligos (20 µM non-template strand DNA, 15 µM template strand DNA, 10 µM RNA, 20 mM Tris-OAc pH 7.7, 5 mM Mg(OAc)_2_, 40 mM KOAc) in a thermal cycler (95 °C for 2 min, 75 °C for 2 min; 45-to-25 °C gradient over 25 min). To reconstitute CTR ePECs, the CTR core RNAP was incubated with the scaffold at 1:2 ratio (2.5 µM CTR-RNAP, 5 µM scaffold) in 10 µL of Elongation Buffer (EB; 25 mM HEPES-KOH pH 8.0, 130 mM KCl, 5 mM MgCl_2_, 0.15 mM EDTA, 5% v/v glycerol, 25 µg acetylated BSA/ml) for 20 min at 37 °C. Reconstitution yielded ≥90% conversion of RNAP to ePECs.

For PAGE analysis, ∼0.3 µL samples were mixed with NuPAGE 4X LDS Sample Loading Buffer (Novex, reducing agents omitted) and electrophoresed through a 4-15% gradient polyacrylamide gel (GE Healthcare) using a PhastSystem Electrophoresis unit (Pharmacia– GE Healthcare). Gels were stained with Imperial Protein Stain (Thermo Scientific), destained with water, and imaged with a CCD camera (Protein Simple).

We used the Fiji software package based on ImageJ to quantify the percentage of crosslinking. The intensity of two crosslinked bands (I_ϕ3-ϕ3′_ for SI3–SI1crosslink, I_ϕ3′-ϕ3′_ SI3–RH crosslink) and the uncrosslinked β and β′ bands (I_unxlink_) were measured. Because the SI3–SI1 crosslink was inter-subunit and the SI3–RH crosslink was intra-subunit, the two crosslink bands were easily distinguished (since MW_ϕ3_=151 kDa and MW_ϕ3′_=155 kDa, we assumed Coomassie brilliant blue G-250 binds β and β′ equivalently). We calculated the % crosslinking from the band intensities (I) in the gel images as:

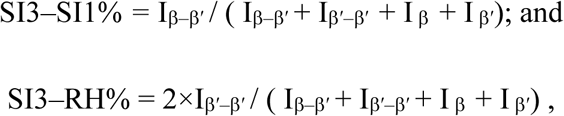

where the crosslinked species are indicated by subscripts containing two subunits and the intensity of the SI3–RH crosslinked species is doubled because it is approximately half the size of the SI3–SI1 crosslinked species. The SPB value was then calculated as the ratio between the two crosslink species:

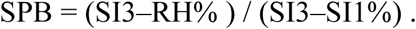

All SPB values were determined from at least three independent assays and reported as mean ± SD.

### Cryo-EM sample preparation

Core RNAP was thawed from storage and subjected to gel filtration (Cytiva Superose 6 Increase, 24ml, 0.5ml/min, 4°C) in buffer L (20 mM TrisCl pH8/RT, 150 mM potassium glutamate, 2 mM MgCl_2_, 5 mM DTT). Peak fractions containing core RNAP were pooled. Template DNA strand (#15254; Key Resource Table) was annealed to 2x molar excess of RNA strand (#15147; Key Resource Table) in annealing buffer (10 mM HEPES pH 7.5/RT, 50 mM NaCl, 1 mM EDTA) by heating to 95 °C for 1 min and passive equilibration to room temperature. Gel filtrated core RNAP was mixed with a 1.3x molar excess of DNA–RNA hybrid scaffold and incubated for 15 min at 37°C. Non-template strand (#15252; Key Resource Table) was added to 1.5x molar excess (with respect to core RNAP concentration) and the sample was incubated for 10 min at 37°C. The complex was concentrated to about 15 μM using Amicon Ultra centrifugal filters (Merck Millipore). Detergent working solutions containing 80 mM 3-([3-cholamidopropyl-]dimethylammonio)-2-hydroxy-1-propanesulfonate (CHAPSO, Sigma #C4695) and 30 mM MgCl_2_ were prepared in buffer L with or without 2 mM ppGpp. The sample and detergent working solutions were centrifuged at 14,000 × *g* for 10 min at 23 °C and the supernatant was transferred to fresh tubes.

### Grid preparation

Sample application and vitrification was performed using a Vitrobot Mark IV (ThermoFisher Scientific) equilibrated to 10°C and 100% relative humidity in the blotting chamber. C-flat holey carbon grids (CF-1.2/1.3-4Au, EMS) were glow-discharged before use. For each condition (+/− ppGpp), 6 μL detergent working solution were added to 58 μL sample and the mix was incubated for 1 min at room temperature. For each grid, 3 μL sample were applied to a glow-discharged grid mounted in the blotting chamber of the Vitrobot instrument. Blotting (blot time 3-4 s, blot force +/−1) and plunging into liquid ethane was performed at once. Approximate final concentrations on the grid were 15 μM complex, 8 mM CHAPSO, and 200 μM ppGpp.

### Nascent RNA extraction

Nascent RNA was purified as described previously^39^ with slight modifications. Frozen cells harvested from 0.5 L cell culture were combined with frozen lysis buffer (20 mM Tris-HCl, pH 8.0, 0.4% Triton-X 100, 0.1% NP-40), 100 mM NH4Cl, 10 mM MnCl_2_, 1x EDTA-free cOmplete Mini protease inhibitor cocktail (Roche)), and 50U/mL RNasin [Promega, N211B]). The mixture was subjected to six rounds of milling at 15 Hz for 3 min in a Retsch MM400 Mixer Mill using 50 ml canisters pre-cooled in liquid nitrogen and 25 mm stainless steel balls. The lysate volume was adjusted to 5.5 mL of lysis buffer by gentle addition of cold lysis buffer and RQ1 DNase I (Promega, 0.054 U/mL) was added followed by incubation for 20 min on ice. The reaction was quenched with EDTA (25 mM final) to release ribosomes from the transcripts and chelate Mg^2+^ to inactivate RNAP. The lysate was then clarified at 4 °C by centrifugation at 25,000 ×g for 10 min and incubated with 0.5 mL anti-FLAG M2 affinity resin (Sigma-Aldrich) for 3 h at 4 °C to isolate RNAP-nascent transcript complexes. The resin was washed four times with lysis buffer with 100 mM NH_4_Cl, 300 mM KCl, 1 mM EDTA, and 50 U/mL RNasin. ECs were eluted with 600 µL lysis buffer with 2 mg/mL FLAG peptide (Sigma-Aldrich), 100mM NH_4_Cl, 1mM EDTA, and 50U/mL RNasin. Nascent RNA was extracted from the eluate with miRNeasy Mini Kit (Qiagen) following manufacturer’s instructions. The nascent RNA sample was further purified by an additional overnight isopropanol-GlycoBlue precipitation at −20 °C, dissolved in DEPC-treated water, and stored at –80°C.

### NET-Seq sample preparation and processing

NET-seq library preparation was performed as described previously^39^ with minor changes to use custom adaptors compatible with sequencing using Illumina NovaSeq 6000. An adaptor oligo (14047) was adenylated using a NEB 5’ DNA Adenylation kit (by incubating 6 µM of the oligo with 80 µM ATP, 6 µM *Mth* RNA ligase (NEB), and 25% (v/v) PEG 8000 in 1X NEB Buffer 1 at 65 °C for 4 hours, inactivated at 85 °C for 5 min, and then and precipitated overnight at −20°C with isopropanol. Precipitated nascent RNA (3 µg total per sample) was then ligated in 3 separate 1 µg reactions to the 5′ adenylated adaptor oligo using a T4 RNA Ligase 2, truncated (T4 Rnl2tr) kit (NEB) with 10% DMSO, 22% PEG8000, 3 µM adenylated DNA linker, T4 Rnl2tr [14.7U/µL], RNasin [2U/µL], and 1x T4 RNA Ligase Reaction Buffer). These ligation reactions were incubated for 4 h at 37°C and the RNA ligase was inactivated with Proteinase K (0.04U/µL) (NEB, P8107). The ligated material was then subjected to cDNA synthesis (1 h) and cDNA circularization reactions (3 h) using CircLigase (Lucigen) as described previously. ^39,47^ Circularized cDNA libraries were PCR amplified as previously described ^47^ with Phusion DNA polymerase (NEB) using PCR conditions recommended by the manufacturer and dual-indexed primers for the NovaSeq6000 (Illumina) sequencing platform. Library concentration and size distribution were determined using an Agilent TapeStation 4150 before 150-nt paired-end sequencing on an Illumina NovaSeq 6000 instrument (sequencing performed by the University of Wisconsin-Madison Biotechnology Center).

### ChIP-seq sample preparation and processing

ChIP-seq was performed as previously described. ^52,86^ A cell pellet corresponding to 50 mL of crosslinked *E. coli* culture was mixed with separately formaldehyde-crosslinked *B. subtilis* RL3116 cells (using a volume of *B. subtilis* cells equivalent to 1% of the *E. coli* cells) and used for each IP reaction. Immunoprecipitation was performed as described ^86^ using an antibody against the beta subunit of RNAP or the FLAG tag on the C-terminal end of the beta prime subunit. Paired-end libraries were prepared using NEBNext Ultra II DNA library reagents according to the manufacturer’s protocols and sequenced on Illumina NovaSeq 6000.

### Intracellular GTP concentration measurement

Metabolites were collected from strains RL3939–3942 by rapid filtration of 5 mL or 25 mL of cells through a circular 47 mm diameter, 0.45 µm pore size nylon filter (Millipore Sigma). The filter with cells was quickly plunged into 3 mL of ice-cold extraction solvent (acetonitrile:methanol:water, 2:2:1). After incubation for ∼10 min on dry ice, the collection filter was flushed with the 3 mL extract and removed. Aliquots (650 µL) of the extracted metabolites were frozen at –80 °C. The samples were filtered, plunged into ice-cold extraction solvent, and stored at –80 °C. To quantify intracellular GTP, 450 µL of metabolite extract was dried under a nitrogen flow, and dried samples were resuspended in 100 µL of solvent A (water:methanol [97:3], 10 mM tributylamine [TBA], 9 mM acetate, pH 8.2) and then analyzed on a Thermo-Fisher Vanquish Horizon UHPLC instrument coupled to a heated electrospray ionization source (HESI) and hybrid quadrupole-Orbitrap high resolution mass spectrometer (Q Exactive or Q Exactive plus Orbitrap, ThermoScientific). Chromatography was performed using a 100 mm × 2.1 mm × 1.7 µm bridged ethylene hybrid (BEH) C18 column (ACQUITY) at 30 °C. Samples (25 µL) were injected onto the column using an autosampler at 4 °C and 200 µL/min flow rate. Small molecules were eluted using mixtures of solvent A and solvent B (100% methanol without TBA) using a 95:5 A:B for 2.5 minutes, followed by a gradient of 90:10 A:B to 5:95 A:B over 14.5 minutes, and then held at a 10:90 A:B for an additional 2.5 minutes. The column was re-equilibrated to 95:5 A:B mixture composition for 5 min between samples. Small molecule data were collected between 3.3 and 18 min and analyzed using mass spec parameters: negative mode scan; 100–800 m/z scan range; 10^6^ automatic gain control (AGC); 3.0 kV spray voltage; 40 ms maximum ion collection time; and 350 °C capillary temperature.

MAVEN (v2.0.1)^87^ was used both to identify peaks corresponding to GTP and to determine areas under identified chromatography peak traces. GTP peaks were gated using the mass-to-charge (m/z) ratio (+/- 25 ppm mass tolerance) pre-programmed for GTP in MAVEN. Peak areas were determined using the MAVEN manual integration–Gaussian smoothing function, which corrected for baseline noise (see Table S10). Peak areas for known molar amounts of GTP (i.e., from the GTP standards) were log-transformed and fit to a linear equation versus log-transformed molar amounts of GTP to obtain a standard curve. Moles of GTP in each sample were determined by interpolation of the log-transformed peak areas using this standard curve. To determine the number of moles of GTP per CFU, the number of moles of GTP was divided by the number of CFUs used to generate the sample loaded onto the C18 column (by converting the OD_600_ to CFU ratio using the qNET-seq spike-in determination described in ‘Determination of qNET-seq spike-in scaling factor’. Moles of GTP per CFU was divided by the estimated volume of a single cell, 7.0 10^−16^ L to give the intracellular GTP concentration (see Table S10).

### Cryo-EM data acquisition and processing

Grids of con-ePEC+ppGpp were imaged using a 300 kV Titan Krios instrument (ThermoFisher Scientific) equipped with a K3 camera (Gatan) and Cs corrector optics. Images were recorded using Leginon with a pixel size of 1.076 Å/px at the detector over a nominal defocus range of −0.8 μm to −2.5 μm. A total of 6,913 movies (“+ppGpp” dataset) were recorded in counting mode (K3 camera binning 1; image dimensions 5,760 x 4,092) with an electron dose of 26 e^−^/ Å^2^/s in dose-fractionation mode of 0.05 s over a 2 s exposure (40 frames) resulting in a total dose of 52 e^−^/Å^2^. Grids of con-ePEC in the absence of ppGpp were imaged using a 300 kV Titan Krios instrument (ThermoFisher Scientific) equipped with a K3 camera (Gatan) and a BioQuantum imaging filter (width = 20 eV). Images were recorded using Leginon with a pixel size of 1.083 Å/px at the detector over a nominal defocus range of −1 μm to −3 μm. A total of 11,885 movies (“−ppGpp” dataset) were recorded in counting mode (K3 camera binning 1; image dimensions 5,760 x 4,092) with an electron dose of 25 e^−^/Å^2^/s in dose-fractionation mode of 0.05 s over a 2 s exposure (40 frames) resulting in a total dose of 50 e^−^/Å^2^. For all datasets, dose-fractionated movies were drift-corrected, summed, and dose-weighted using MotionCor2.^76^ The contrast transfer function (CTF) was estimated for each summed image using the Patch CTF module in cryoSPARC v3.3.^88^ Micrographs were curated to remove outliers in estimated CTF fit resolution resulting in 6,536 images for the +ppGpp dataset (5.4% rejected) and 11,698 images for the −ppGpp dataset (1.6% rejected). Particles were picked using cryoSPARC Blob Picker (circular blob, 150 Å < x < 200 Å, minimum separation 0.5) and extracted with a box size of 384 px. Extracted particle images were subjected to three rounds of cryoSPARC 2D classification (150-300 classes, batch size 400, uncertainty factor = 4) before generating ab initio reconstructions in cryoSPARC^88^ yielding maps of RNAP or “junk”. Particles were further curated using two rounds of cryoSPARC Heterogeneous Refinement (N=6) with ab initio classes serving as 3D references (2 RNAP and 4 junk references). Particles contributing to RNAP classes were combined and curated for duplicates (100 Å center-center distance cutoff). 3D reconstructions were refined using cryoSPARC Non-Uniform (NU) Refinement with Defocus and Global CTF Refinement (tilt/trefoil) enabled^74^ yielding consensus reconstructions nominal resolution values of 3.1 Å (597,682 particles) for the +ppGpp dataset and 2.7 Å (1,257,951 particles) for the −ppGpp dataset. Particle sets were subjected to RELION v3.1 Bayesian Polishing.^75^ Per-particle motion-corrected particles were imported into cryoSPARC and subjected to NU Refinement with Defocus and Global CTF Refinement (tilt/trefoil) enabled, yielding a consensus reconstruction with a nominal resolution of 2.8 Å for the +ppGpp dataset and 2.5 Å for the −ppGpp dataset. Particles were further curated using RELION v3.1 masked 3D classification with signal subtraction using mask “omega” to remove particles without the omega subunit (16.7% removed in +ppGpp dataset; 25.2% removed in −ppGpp dataset), followed by removal of particles containing Sig70 (6% removed for +ppGpp dataset; 9% removed for −ppGpp dataset) using mask “sigma”.

To resolve heterogeneity of functional states, the particle stack of the +ppGpp dataset (468,267 particles) was further subjected to RELION v3.1 3D classification with signal subtraction using a mask encompassing the RNAP main channel (mask “NA”). Classification yielded a pre-translocated (180,983 particles) and a half-translocated state (243,833 particles). 3D Variability Analysis (3DVA) in cryoSPARC (3 components) was performed on signal subtracted particles of each state using a mask encompassing the swivel module, the active site, and the downstream channel (mask “swivel”). The half-translocated state exhibited no conformational heterogeneity and was considered a final class (243,883 particles; 2.8 Å). 3DVA of the pre-translocated state revealed large conformational changes of the SI3 domain and rim helices (Supplementary Videos 1-3). Subsequent clustering (N=4) and pooling according to functional state yielded three classes: fTL-FL/RH down (23,591 particles; 3.2 Å), fTL-FL/RH up (53,279 particles; 3.1 Å), and uTL (104,112 particles; 3.0 Å).

Functional heterogeneity of the −ppGpp dataset was resolved following analogous steps. The 7-containing, ο−^70^-less particle stack was subjected to RELION v3.1 3D classification with signal subtraction using a mask encompassing the RNAP main channel (mask “NA”).

Classification yielded one class for a pre-translocated state (203,144 particles), one class for a half-translocated state (215,551 particles), and several classes exhibiting a poorly defined translocation state (termed “intermediate”; 420,818 particles). 3D Variability Analysis (3DVA) in cryoSPARC (3 components) was performed on signal subtracted particles of each state using a mask encompassing the swivel module (mask “swivel”), the active site and the downstream channel. 3DVA and subsequent clustering (N=8) of the intermediate translocation state particle stack yielded a vast conformational heterogeneity of the swivel module (Supplementary Videos 4-6). Two clusters exhibited a half-translocated state and were pooled with the main stack of half-translocated particles. The remaining clusters were pooled into a single “partially swiveled” class (314,501 particles; 2.7 Å). The pooled half-translocated exhibited no conformational heterogeneity and was considered a final class (321,868 particles; 2.6 Å). 3DVA of the pre-translocated state revealed large conformational changes of the SI3 domain and rim helices (Supplementary Videos 7-9). Subsequent clustering (N=4) and pooling according to functional state yielded three classes: fTL-FL/RH down (65,808 particles; 2.8 Å), fTL-FL/RH up (73,863 particles; 2.8 Å), and uTL (63,437 particles; 2.8 Å).

Local resolution and locally filtered maps were generated using blocres/blocfilt.^77^ Most structural biology software was accessed through the SBGrid software package. ^73^

### Model building and refinement

Initial models were derived from PDB 8EFH, 8EGB, 8EHA, and 8EG8.^22^ The starting model was manually fit into the cryo-EM density maps using ChimeraX^78^ and further modified using ISOLDE^79^ and Coot.^80^ The model was finalized by all-atom and B-factor refinement with Ramachandran and secondary structure restraints using phenix.real_space_refine against the full/unsharpened map.

Maps and models were visualized in ChimeraX.^78^

### Determination of swivel module rotation angle

Rotations angles of the swivel module were calculated as described previously using a post-translocated EC (PDB 6ALH)^23^ as a reference.^25^ Structures were first aligned using the following

PyMOL command.

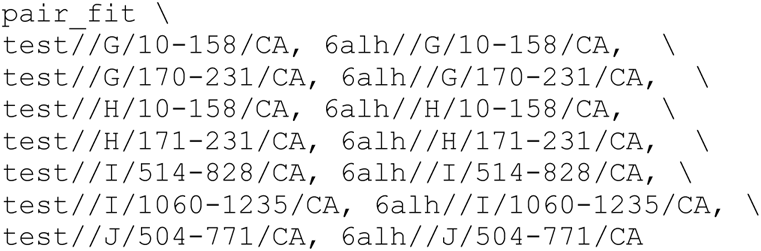

The swivel domain was defined using the following PyMOL command:

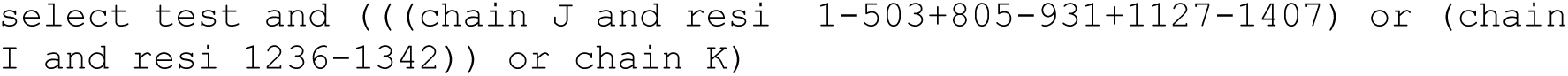

### NET-seq data processing and scaling

NET-seq data were processed using standard tools and scaled for quantitative comparison across samples based on the number of *B. subtilis* spike-in reads that mapped uniquely to *B. subtilis* genome. Adapters, linker, and control oligos potentially contaminating each sample were trimmed from raw reads using cutadapt (v3.4).^82^ Reads with a minimum length of 15 nt were mapped to the MG1655 genome (NC_000913.3) using Bowtie (v1.3.0) allowing one mismatch and allowing random assignment of reads mapping to multiple loci based on alignment stratum (Bowtie options --best -a -M 1 -v 1).^83^ Alignments for all samples, except those derived from *E. coli* RL3938, were converted to BAM and BED files using SAMtools (v1.16.1) ^84^ and BEDtools (v2.30.0).^85^ To bring mapped reads from RL3938 into parity with those of the other strains, alignments were randomly down-sampled to 30% of mapped reads using samtools (option -s 10.3) prior to conversion to BED files. The specific 3′-end counts for each genome position were determined using bedtools (options -d -strand - -5 [plus strand] or -d -strand + -5 [minus strand]). To scale read count between samples, the 3′-end counts mapped to each genome position were scaled to total read counts per ten million and multiplied by the *B. subtilis*-unique scaling factor (Table S2).

### Determination of qNET-seq spike-in scaling factor

To determine the spike-in scaling factor for each sample, we first identified specific regions in the *B. subtilis* genome (Accession: NC_000964.3) to which *E. coli* reads did not map (*Bsu* unique regions) using an *E. coli-*only NET-seq dataset. We mapped reads from the *E. coli*-only NET-seq library to a combined *E. coli* MG1655 and *B. subtilis* strain 168 genome file using Bowtie with options to suppress reads that map to multiple loci and allowing no mismatches (options –best -a -m 1 -v 0). After conversion to BAM and BED files, total locus coverage (read coverage not 3′-end coverage) was determined for the *B. subtilis* strain 168 genome. We then generated a BED table of regions with zero coverage (i.e., regions of the *B. subtilis* genome to which *E. coli*-only reads did not map) to detect *B. subtilis-*unique reads present in *E. coli* NET-seq samples containing a *B. subtilis* spike-in.

To determine the number of *B. subtilis*-unique reads in each sample with *B. subtilis* spike-ins, we followed the same NET-seq processing pipeline used to generate the table of *B. subtilis* unique loci above but used Bedtools to count 3′-end coverage for both *B. subtilis* unique reads and reads mapping to both *B. subtilis* and *E. coli* (i.e., not mapping to the *B. subtilis* unique loci). To determine the *B. subtilis*-unique scaling factor for each sample, the fraction of *B. subtilis*-unique reads was calculated as the *B. subtilis*-unique counts divided by the total number of reads mapping to *E. coli* and *B. subtilis* loci (Table S2). The smallest *B. subtilis*-unique fraction was identified among the set of samples to be compared, and this number was divided by the *B. subtilis*-unique fraction for each sample to obtain the *B. subtilis*-unique scaling fraction. The scaling fraction was converted to a scaling factor by multiplying it by 10^7^ divided by the total reads mapped to *E. coli*. The scaling factor was then used to generate scaled reads per 10^7^ reads for each sample. For most analyses, the average of the two scaled reads per genome position was used.

### Pause calling from qNET-seq reads

No standard criteria to identify pause sites from NET-seq data exists; various approaches have been used in studies to date.^31^ ^Gajos, 2021 #109,38,39^ Here, we modified the original method of Larson et al. (z>4 in a 200 bp window) to improve identification of closely spaced pauses by adjusting the mean used to calculate *z* scores to reduce the masking effect of closely spaced pauses on *z*. Pauses (Tables S3,S4,S11–S14) were identified using a custom perl script (pause_ratio_by_position.pl) to analyze averaged and scaled qNET-seq reads for pairs of isogenic strains containing pALS13 (RelA*, high ppGpp) or pALS14 (RelA^−^ control, basal ppGpp). Pause sites were assigned to locations in each strain in which the read counts was ≥4 standard deviations from the adjusted mean of read counts for the surrounding +/– 100-bp window (i.e., adjusted *z* ≥ 4). To reduce cases where closely spaced pauses might be missed by their effect on the mean and to allow identification of up to three pauses within each 201-bp window, the three highest read positions were omitted when calculating the adjusted means used to determine if the counts at a given position met the *z* ≥ 4 criterion. Pause strength (PS) was calculated as the ratio of the read counts at a position to the adjusted mean in the surrounding +/–100-bp window.

### Calculation of qNET-seq and pausing parameters

qNET-seq occupancies (Figures 4B, 5B, 5F, S4A, and S4B) were calculated as the sum of scaled qNET-seq counts per unit DNA (e.g., kb). Relative occupancies were assigned per segment of DNA (Figure 4B) or were calculated as rolling averages in 201-bp windows (Figures 5B & 5H). Total sense versus antisense occupancies (Figure S5A) were calculated as the proportions of sense and antisense counts for all genes. The distributions of antisense/sense occupancy ratios (Figure S5A) were calculated for the 1000 genes >500 bp in length with highest sense occupancy. Average qNET-seq pause frequencies (Fig. 4C) were calculated for sets of expressed genes >1 kb in length at the 5′ ends of transcription units in 21 bp windows for the first 1 kb of the genes. Expressed genes were determined from the qNET-seq occupancies of the genes with a cutoff selected to give the same numbers of genes for datasets being compared (e.g., 350 counts/kb yielding the two-thirds most highly expressed genes (∼680 genes) for Figure 4C (Figure S2). The pause frequencies were converted to pause frequency per 100 bp for plotting.

Aggregate PSs (Figs. 4C & 5G) were calculated for the same sets of genes and converted to average PS/10 bp for plotting.

### ChIP-seq data analysis

ChIP-seq data (Figures 5F, 5I, and 5J) were processed as described previously,^86^ with slight modifications. Read quality control was performed using fastqc (https://www.bioinformatics.babraham.ac.uk/projects/fastqc/) and multiqc. ^89^

All ChIP-seq analyses were performed using an open-source-software-based ChIP-seq pipeline (https://github.com/mikewolfe/ChIPseq_pipeline). The pipeline manages computational jobs using Snakemake^90^ and maintains the relationship between samples and their metadata using peppy.^91^ Paired- and single-end reads were trimmed of adapter sequences using CutAdapt with parameters ‘-a AGATCGGAAGAGCACACGTCTGAACTCCAGTCA -A

AGATCGGAAGAGCGTCGTGTAGGGAAAGAGTGT’. Quality trimming was performed using trimmomatic^92^ with parameters “LEADING:3 TRAILING:3 SLIDINGWINDOW:4:15”. The reads were aligned to the *E. coli* (Genbank accession: U00096.3) and *B. subtilis* (Genbank accession: AL009126.3) chromosomes using bowtie2 (16) with parameters “-- end-to-end --very-sensitive --phred33”. Raw read coverage was calculated using deepTools^93^ with the parameters “--binSize 5 --samFlagInclude 67 --extendReads”. The raw *E. coli* coverage counts were first scaled by multiplying the raw coverage by a spike-in based scaling factor determined as described.^94^ Briefly, a background subtracted fragment count was determined in the spike-in fragments by taking the raw fragment counts across the *B. subtilis* chromosome AL009126.3 for paired extracted and input samples, dividing the read fragments by the total number of *B. subtilis* read fragments per million for each sample and then determining a background subtracted fragment count as follows:

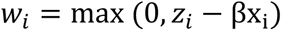

Where, 𝑤_!_ is the background subtracted fragment count per position 𝑖, 𝑧_!_ is the fragment counts in the extracted sample, 𝑥_!_ is the fragment counts in the paired input sample, and β is a regression-based scale factor determined by linear regression with regions outside the full coding region of all genes plus an additional 300 bp upstream of each gene. The average enrichment for a given sample, k, was calculated using the background subtracted fragment count per position j in each coding region plus an additional 300 bp upstream for each gene as follows:

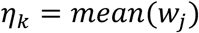

Finally, the spike-in based scaling factor for each sample k was determined as follows:

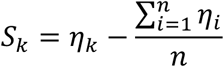

Where n is the total number of samples to be scaled together. This spike-in factor was then applied to the fragments mapping to the *E. coli* genome by taking the raw fragment counts and first scaling by the total number of fragments per million that mapped to the *E. coli* genome per sample and then scaling by the spike-in scale factor 𝑆_#_ described above. After initial scaling, all ChIP traces were copy-number-corrected to account for changes in DNA content between the two genotypes as described in the input correction section below. Occupancies were calculated as an average per position over all defined regions (Figure 5I).

### DNA-input correction

To estimate DNA copy number, DNA was isolated and sequenced before ChIP-seq enrichment. Input libraries were processed identically to the ChIP-seq libraries and a smooth function consisting of a weighted combination of cubic-cyclic basis splines was fit to the raw read coverage separately for each sample using a generalized additive model with the python package statsmodels.^95^ A position-dependent scaling function relative to the input average raw coverage was generated from each model and used to correct for changes in DNA copy number for each ChIP-seq and NET-seq sample.

### Determination of RNAP occupancy on genome features from qNET-seq and qChIP-seq data

To estimate the number of RNAP molecules occupying different genes, gene segments (AUG-proximal vs. AUG-distal), and genome features from qNET-seq and qChIP-seq reads as shown in Figure 5E, 5F, 5H, 5I, and S5, qNET-seq and qCHIP-seq read counts were summed over annotated features of *E. coli* K-12 (NCBI Refseq U000913.3) using custom Perl scripts written for specific analyses (Eco_genome_occupancy_classifier.pl, split_gene_occ.pl, split_gene_chip_occ.pl). For qNET-seq, sense and antisense read counts were summed separately for annotated coding and non-coding genome features (Tables S17 and S18). Total scaled qNET-seq counts were assigned initially to categories: (1) protein-coding genes, (2) antisense to protein coding, (3) tRNA genes, (4) sRNA genes, (5) intergenic regions including REP elements, (6) antisense to tRNA, sRNA, rRNA genes (ncRNA antisense), and (7) other loci, which included pseudogenes, IS elements, and prophage genes. Reads mapping to the rRNA genes and to ncRNA 3′ nucleotides were not included in these assignments because contamination of the qNET-seq counts was readily apparent from the high numbers of counts mapping rRNA genes and tRNA, sRNA, and rRNA 3′ ends. The tRNAs and sRNAs were assumed to be largely undegraded and so only the 3′ end counts were removed, whereas degradation of rRNA was evident causing us to remove internal *rrn* 3′ ends also. tRNA genes from *rrn* operons were added into tRNAs. All other qNET-seq counts were accounted for in these 7 categories. Counts mapping to *rrn* operon leader regions and *rrn* operon spacer regions were summed separately and used to estimate levels of *rrn* operon transcription that were not confounded by rRNA contamination. The spacer regions were used in calculations because qNET-seq can miss counts at near the 5′ ends of transcripts. The rRNA counts were modeled as an eighth category of rRNA operons as described below.

To convert NET-seq read counts to estimates of population-averaged RNAP occupancies on DNA features, counts per RNAP was estimated from counts mapping to the *rrn* operon intergenic regions, the number of RNAPs on *rrn* operons observed by electron microscopy,^96^ the rates at which *rrn* transcription decays after ppGpp induction,^14^ and the copy number of rRNA operons determined from the levels of DNA sequence reads in qChIP-seq input samples (total cell DNA before IP; see DNA Input Correction). Assuming the *ter* region was present at 1 copy per cell, we calculated *ori*-region and *rrn*-operon copy numbers for RNAP^+^ and site 1^−^ cells at basal ppGpp (3.2, ∼18.2, respectively), for RNAP^+^ cells at high ppGpp (2.7, ∼15.8, respectively), and for site 1^−^ cells at high ppGpp (2.2, ∼13.4, respectively). In rapidly dividing cells in rich medium, the *rrn* operons contain 65 ± 2 RNAP molecules.^96^ To determine qNET-seq reads counts arising from nascent *rrn* transcripts present prior to cell harvest, we first summed the counts that mapped to the spacer regions in the *rrn* operons between 16S and 23S *rrn* genes, excluding any tRNA genes present. We then multiplied this count sum by the ratio of total *rrn* operon bp to total spacer bp (37886/2247). Because the level *rrn* transcription decreases upon ppGpp induction,^14^ we used the measured rates of decrease after induction in RNAP^+^ and site 1^−^cells to predict the qNET-seq *rrn* reads that would be present prior to cell harvest. We then divided these total predicted *rrn* transcript counts by the calculated number of transcribing RNAPs (65 per *rrn* operon times the *rrn* operon copy number). This calculation yielded 1182, 868, and 1027 RNAPs, respectively, for basal ppGpp, RNAP^+^–high ppGpp, and site 1^−^–high ppGpp cells. Dividing these numbers into the predicted total *rrn*–operon qNET-seq counts yielded a remarkably consistent number for qNET-seq read counts per transcribing RNAP (2720–2863; average of 2797 counts/RNAP). To obtain the number of qChIP-seq counts per RNAP, we summed qNET-seq and qChIP-seq counts for a set of 540 genes >1 kb that were distal to TSSs and used the ratio of total qChIP-seq counts to total qNET-seq counts to estimate ∼31484 qChIP-seq counts/RNAP. Dividing the total qNET-seq read counts attributable to transcribing RNAPs by 2797 qNET-seq counts/RNAP yielded ∼3200 transcribing RNAPs per RNAP^+^ cell and ∼2750 transcribing RNAPs per site 1^−^ cell. Previous estimates for the number of transcribing RNAPs for *E. coli* growing in rich medium range from ∼1000 to ∼5000,^59,97^ with only the latter including classes other than protein-coding and *rrn* (*e.g*., antisense and intergenic).

Thus, our calculated numbers of transcribing RNAPs are reasonable given the range of prior estimates. We therefore used these estimates to catalog fractions of transcribing RNAPs on different classes of genomic DNA elements (Figures 5E, 5F and S5; Tables S17 and S18). However, we emphasize that these calculations rely on multiple assumptions and thus are primarily useful to compare relative RNAP numbers among genomic element classes in the different strains and conditions we tested.

## Notes

### Competing Interest Statement

The authors have declared no competing interest.

