## Supplementary Figures for "ppGpp regulates transcription elongation via direct and indirect inputs to RNA polymerase pausing and nucleotide addition"

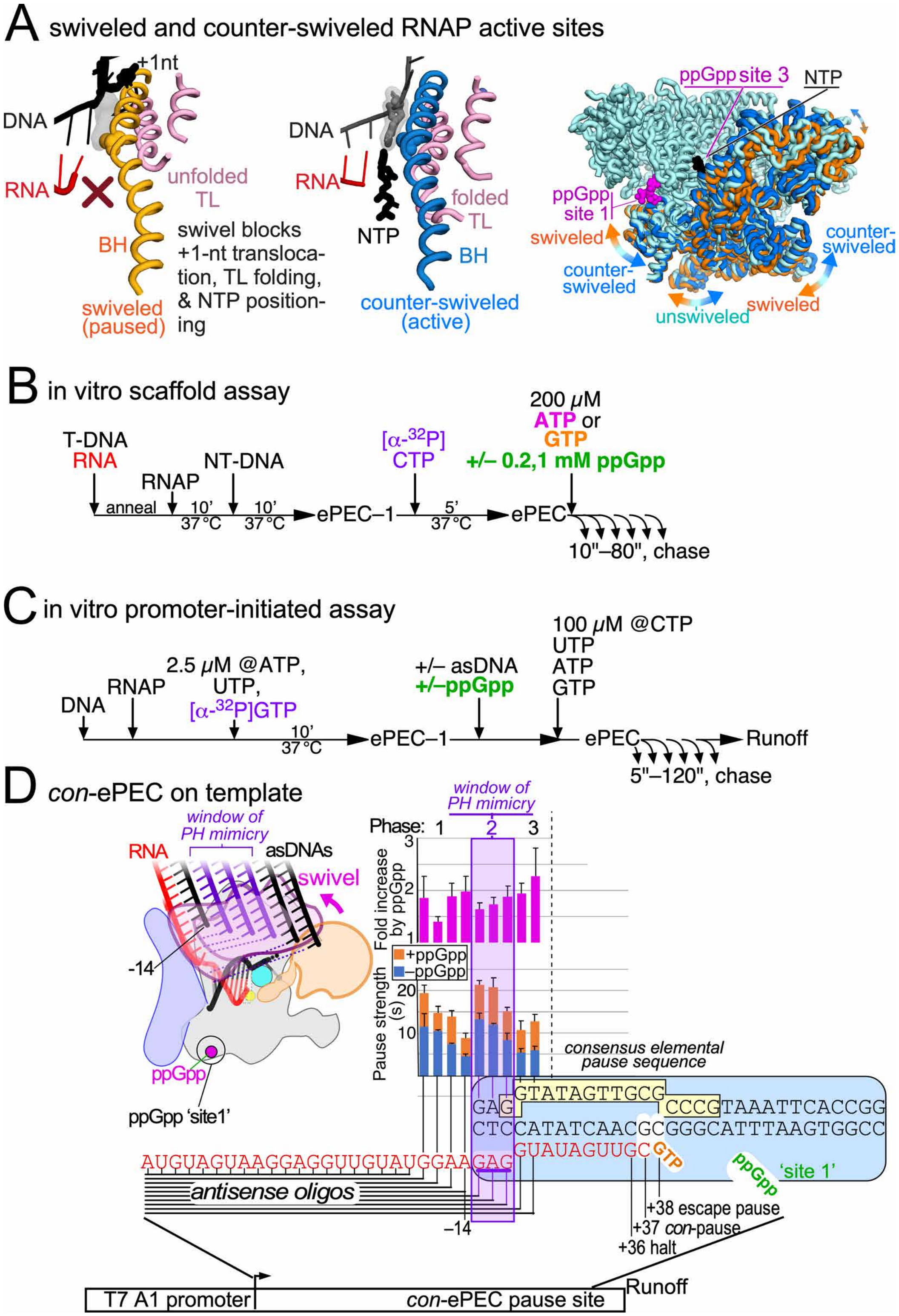


**FIGURE S1.** ppGpp stimulates pausing by affecting swiveling but does not depend on backtracking. Related to Figure 1.

**A.** Swiveling kinks the bridge helix (BH), which inhibits translocation of +1 template DNA and folding the catalytic trigger loop (TL) thereby increasing the lifetime of transcriptional pauses. Conversely, counter-swiveling is associated with TL folding in the presence of bound NTP.
**B.** Experimental scheme to measure pause strengths on synthetic scaffolds.
**C.** Experimental scheme to measure pause strengths on promoter-initiated DNA templates.
**D.** Effects on pausing with and without ppGpp of antisense DNAs (asDNAs) complementary to nascent RNA. asDNAs that block backtracking do not eliminate pause stimulation by ppGpp whether or not they mimic the pause-stimulatory action of a pause hairpin.

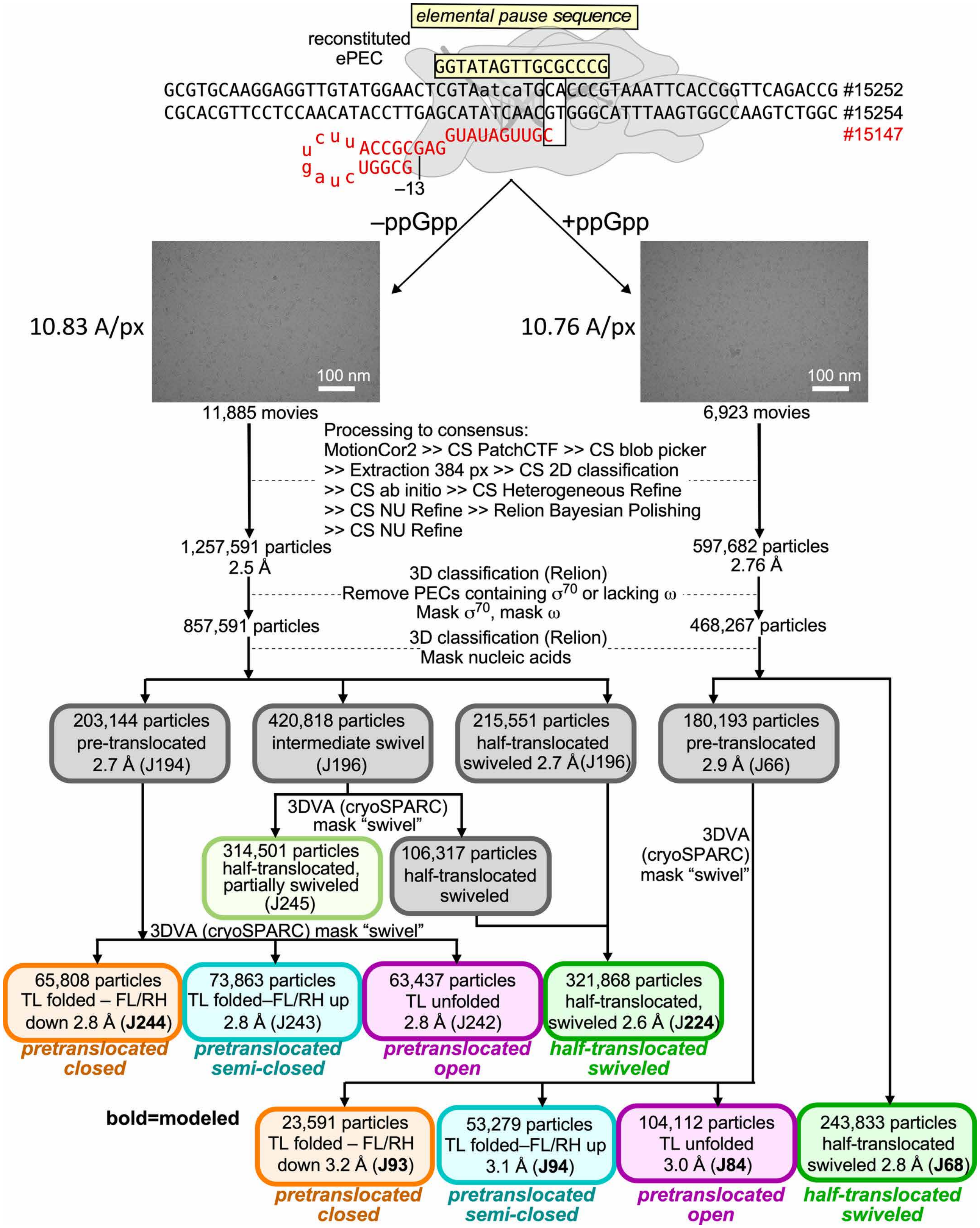


FIGURE S2. Cryo-EM particle sorting. Related to Figure 2

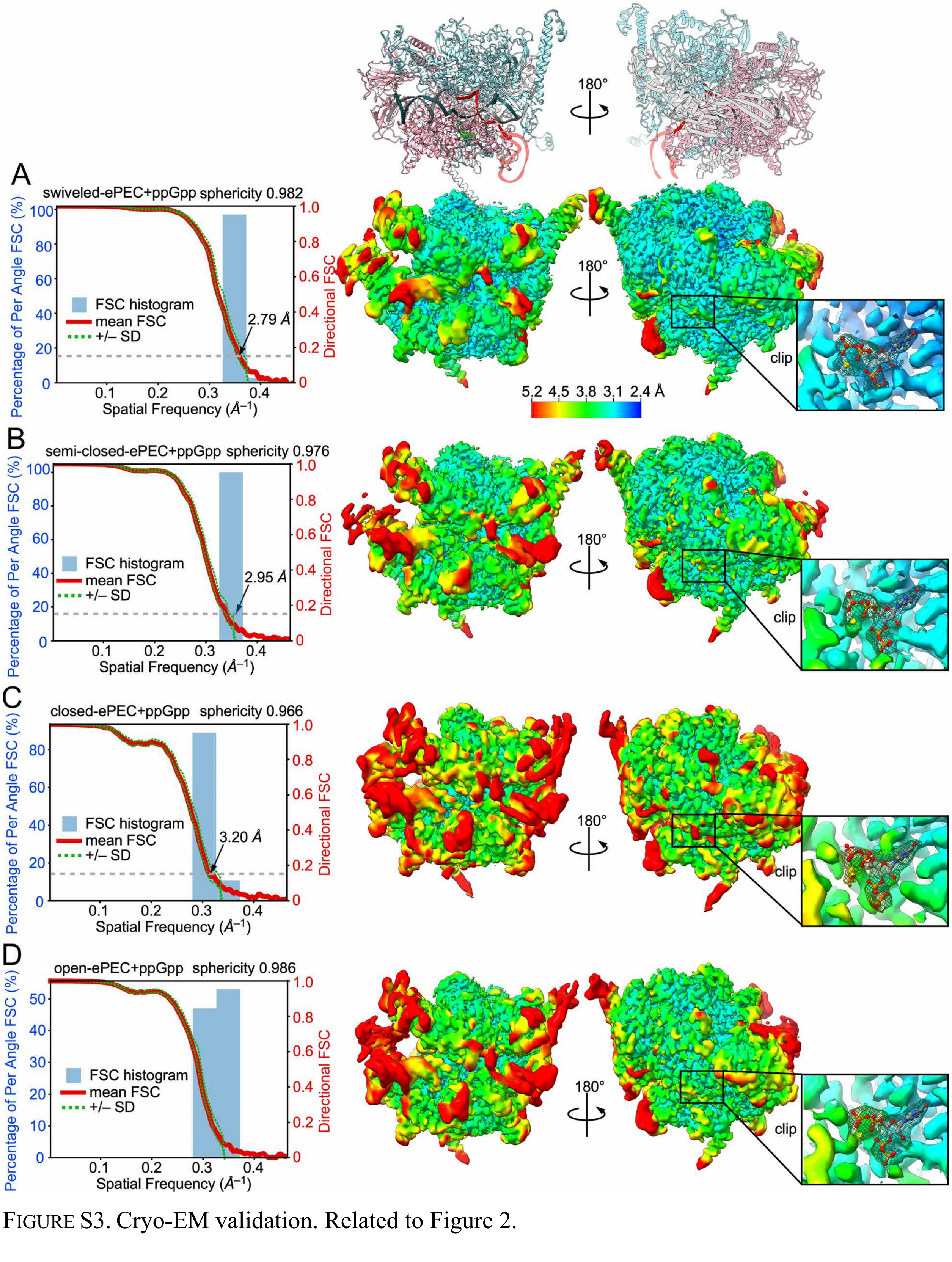


FIGURE S3. Cryo-EM validation. Related to Figure 2.

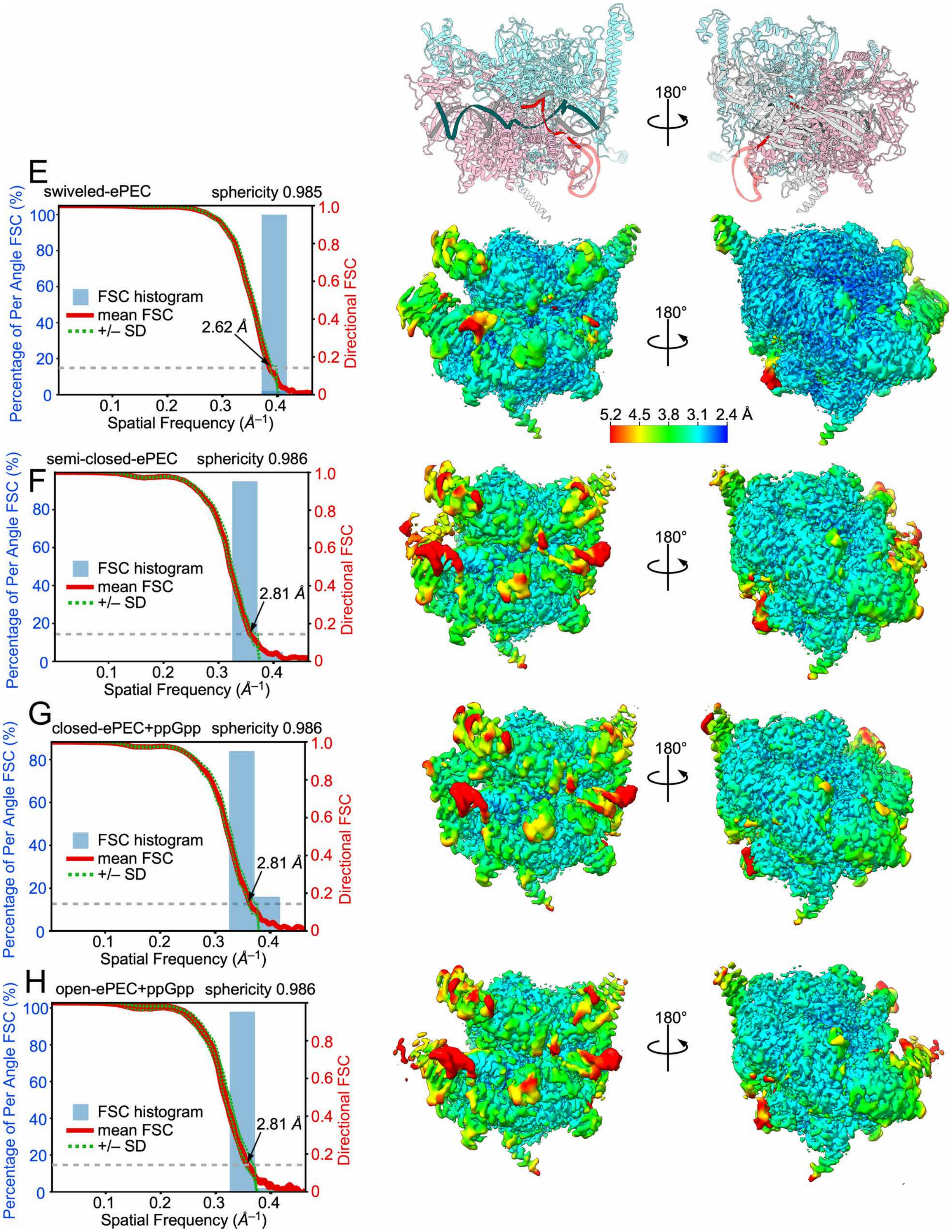


FIGURE S3. Cryo-EM validation. Related to Figure 2, continued.

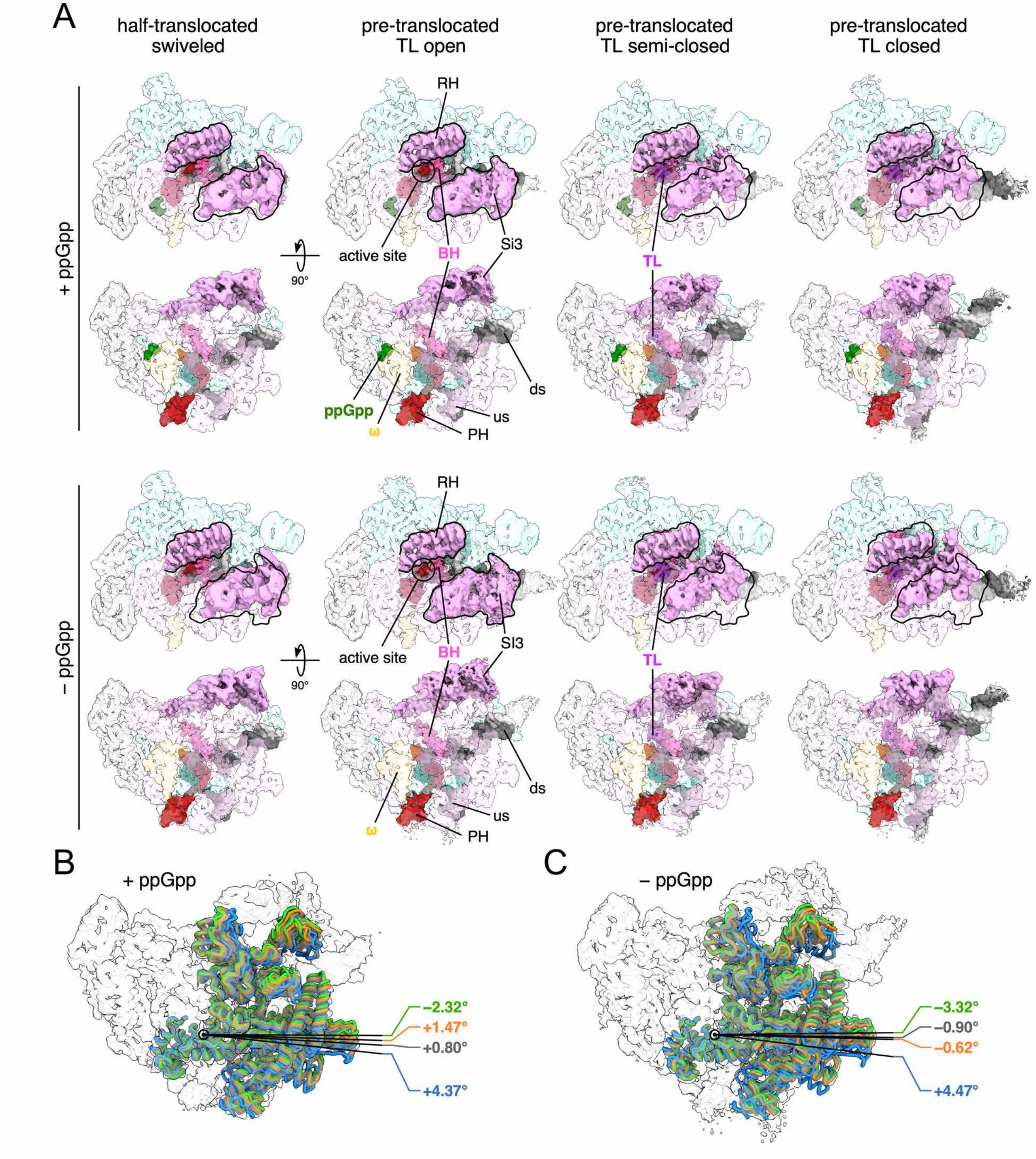


FIGURE S4. RNAP modules in different ePEC conformations. Related to Figure 3.

**A.** Cryo-EM maps of ePEC conformations with and without bound ppGpp. Light pink, rim helices (RH) and SI3. Dark pink, bridge helix (BH). Violet, trigger loop (TL). Green, ppGpp. Red, RNA. Light and dark gray, template and nontemplate DNA, respectively. Black outline, Positions of the RH and SI3 in the open ePEC.

**B,C.** positions of the swivel module with (B) and without (C) bound ppGpp shown as cartoons within the cryo-EM maps for the swiveled (blue), open (orange), semi-closed (gray), and closed (green) ePEC conformation.

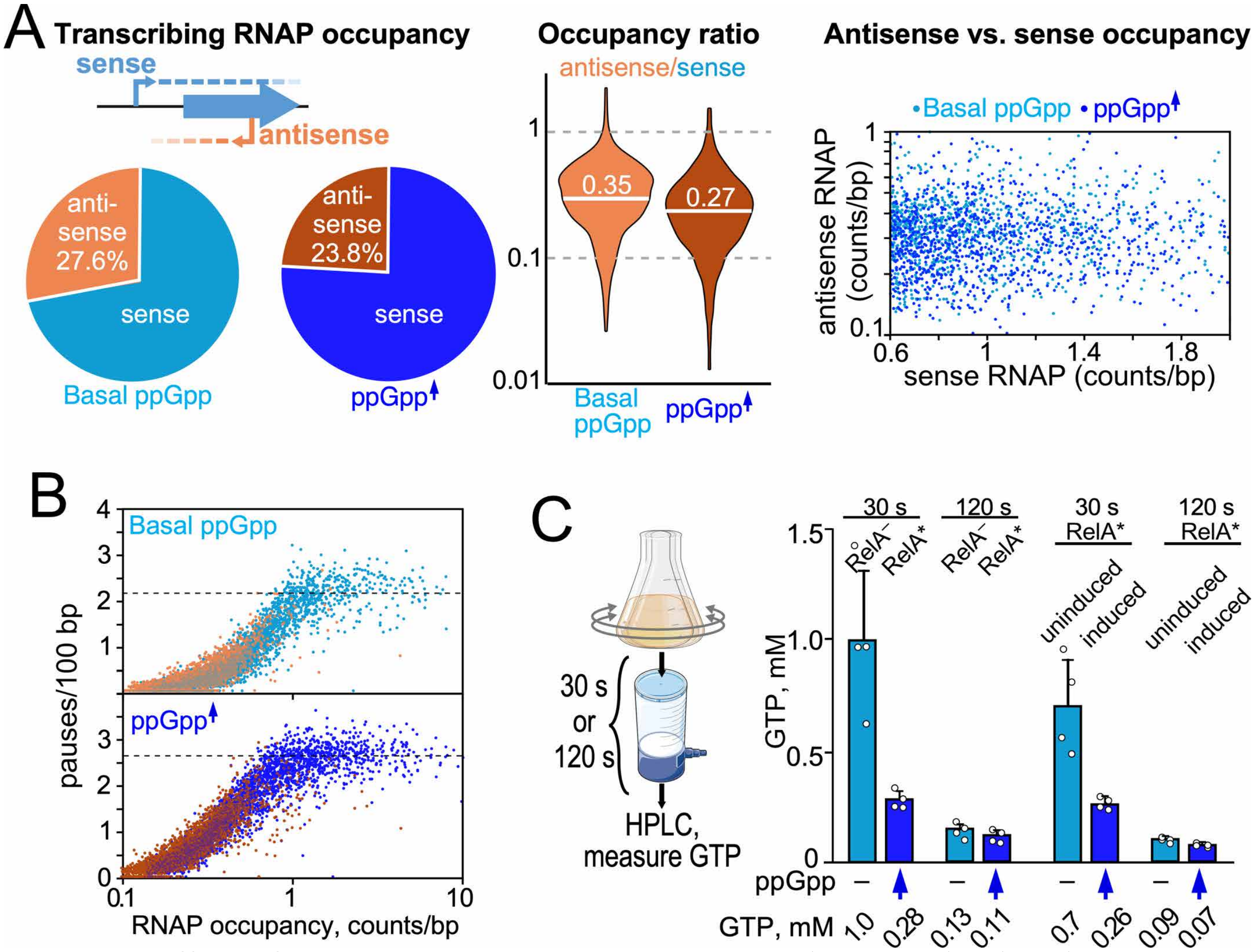


FIGURE S5. Effects of ppGpp on RNAP occupancy, pauses, and GTP. Related to Figure 4.
**A.** Antisense transcription in *E. coli* K-12 assayed by qNET-seq. The pie charts depict the fractions of reads in protein-coding genes mapped to the sense and antisense strands at basal and high ppGpp. The violin plot depicts the ratio of antisense to sense reads in protein-coding genes at basal and high ppGpp. The scatter plot illustrates the lack of correlation between levels of sense and antisense transcription in protein-coding genes as counts/bp in reads mapped to the sense and antisense strands at basal and high ppGpp.
**B.** Pauses/100 bp versus RNAP occupancy (average scaled qNET-seq counts/bp) for basal and high ppGpp conditions for sense (blue) and antisense (orange) transcription. The decrease in mapped pauses at lower occupancy reflects the use of a count cut-off in pause calling (Methods) as well as the lower statistical power to call pauses at low occupancies. The overlap in the sense and antisense distributions indicates that RNAP pausing behavior is similar in sense and antisense transcription.
**C.** Intracellular GTP concentration measured by filtration and extraction for comparison to NET-seq cell harvest filtration. Two technical replicates from each of two independent biological replicates of each strain and condition were measured (Methods). Where indicated, IPTG was added to 1 mM 5 min before filtration of cells.

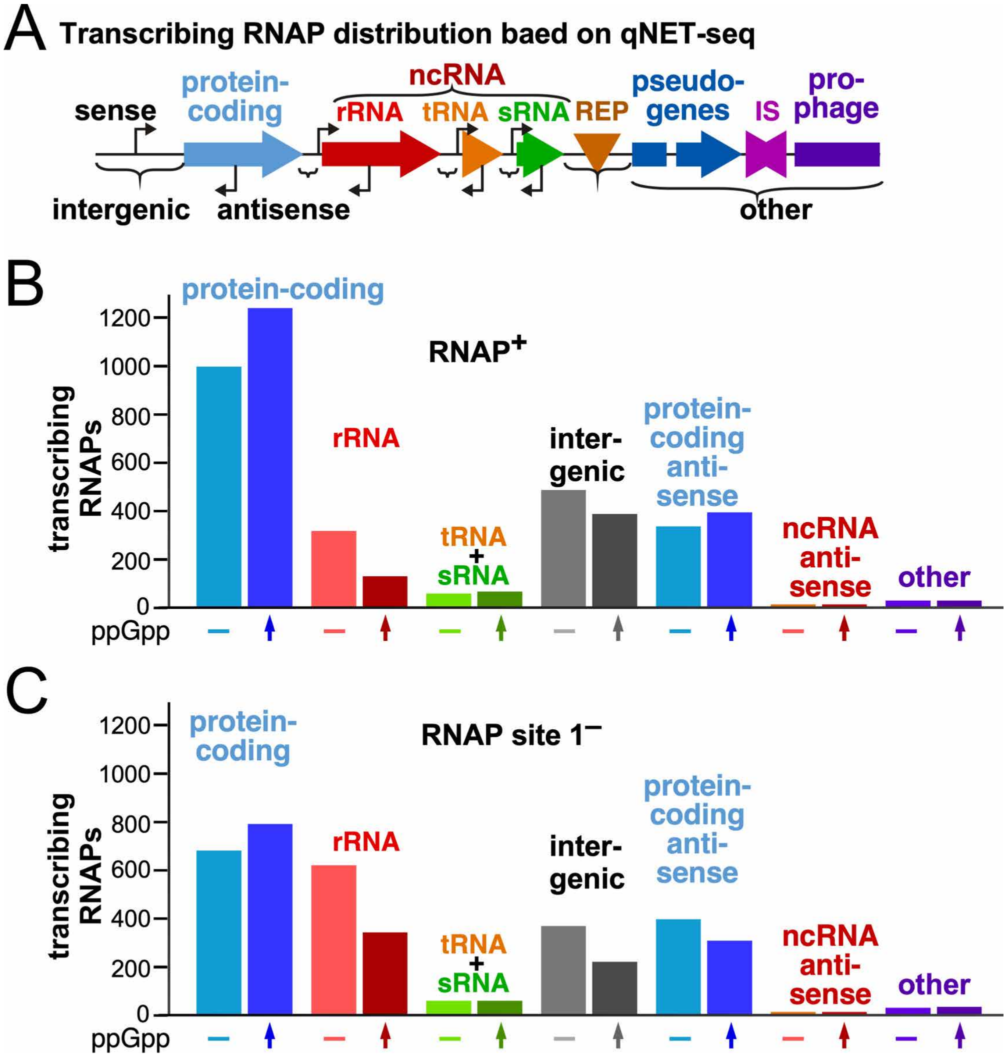


FIGURE S6. Related to Figure 5. RNAP occupancy on feature classes.
**A.** Feature classes.
**B**. Relative numbers of transcribing RNAPs on features per cell.
